# Age-specificity in territory quality and spatial structure in a wild bird population

**DOI:** 10.1101/2024.03.11.584379

**Authors:** Joe P. Woodman, Ella F. Cole, Josh A. Firth, Ben C. Sheldon

## Abstract

Age influences behaviour, survival, and reproduction; hence variation in population age structure can affect population-level processes. The extent of spatial age structure may be important in driving spatially-variable demography, particularly when space-use is linked to reproduction, yet it is not well understood. We use long-term data from a wild bird population to examine spatial age structure and quantify covariance between territory quality and age. We find associations between age and aspects of territory quality, but little evidence for spatial age structure compared to territory quality and reproductive structure. We also report little between-year repeatability of spatial age structure compared to structure in reproductive output. We suggest that high breeding site fidelity and frequent territory turnover by younger breeders, driven by high mortality and immigration rates, limits the association between age and territory quality and weakens overall spatial age structure. Greater spatial structure and repeatability in reproductive output compared to age suggests that habitat quality may be more important in driving spatially-variable demography than age in this system. We suggest that the framework developed here can be used in other taxa to assess spatial age structure, particularly in longer-lived species where we predict from our findings there may be greater structure.

## 1 Introduction

Age affects many aspects of life, as resources are allocated into processes and traits at different points throughout lifespan to maximise fitness (Stearns 1992) and individuals gain in experience as they age. Further, variation in population age structure may affect population dynamics, via effects on reproductive output and social organisation (Coulson et al. 2005; Gamelon et al. 2019; Siracusa et al. 2023; Woodman et al. 2024). There is widespread evidence of temporal variation in age structure in wild populations (Coulson et al. 2005; Gamelon et al. 2016; Hoy et al. 2020), but the way in which age structure varies across space seems to be much less well understood. Some work identifies spatial variation in age structure in fish populations, often linked to human harvesting of different age-cohorts in given areas (Fowler et al. 2000; McIlwain et al. 2005); and in ungulate and bird populations, associated with age-specificity in habitat use and social behaviour (Ferrer and Bisson 2003; Albery et al. 2022). Given that age is often important for individuals’ fitness, spatial age structure is particularly interesting as it might drive local age-related population dynamics. For example, if there is covariance between habitat quality and the age of individuals occupying sites, both could lead to spatially-variable demography, thus potentially biassing estimates of the effects of age on reproduction or, conversely, estimates of environmental effects on trait variation.

One social system where covariance between age and habitat might lead to local age structure is where individuals defend breeding territories. Here, age structure might be particularly important in driving spatial age-related dynamics due to the tight association between breeding territories and reproductive output (Kerbiriou et al. 2006; Jones et al. 2014; Groenendijk et al. 2015). In many animal taxa, individuals undergo reproductive attempts while defending a territory in which resources are used for breeding (Reynolds 1996; Davies et al. 2012). This is particularly prevalent in socially monogamous bird species (Hinde 1956; Greenwood 1980). In such cases, spatial age structure could develop as non-randomness arises when individuals of either similar or dissimilar age breed in closer proximity to each other than expected from chance.

Several mechanisms might affect spatial age structure in territorial animals; we outline four mechanisms here. First, temporal variation in population age structure might influence how age is arranged in space. Temporal variation in wild population age structure is common, particularly in short-lived species where the proportion of populations consisting of the youngest age-cohorts may vary greatly between breeding seasons (Gamelon et al. 2016; Woodman et al. 2022). Such variation might affect how age is arranged in space through passive mechanisms whereby clusters of territories occupied by same-age individuals will be more likely to arise when the age distribution is skewed towards this age-cohort. For example, recent work demonstrates how fluctuations in population age structure determine how age is structured within breeding pairs, where there is greater age-assortative pairing when the proportion of yearlings is higher (Woodman et al. 2022). Thus, between-year age distribution is likely to passively affect spatial age structure as clusters of territories occupied by same-age individuals are more likely to arise when much of the population exists in a single age-cohort (Fig. 1a).

**Fig. 1.**
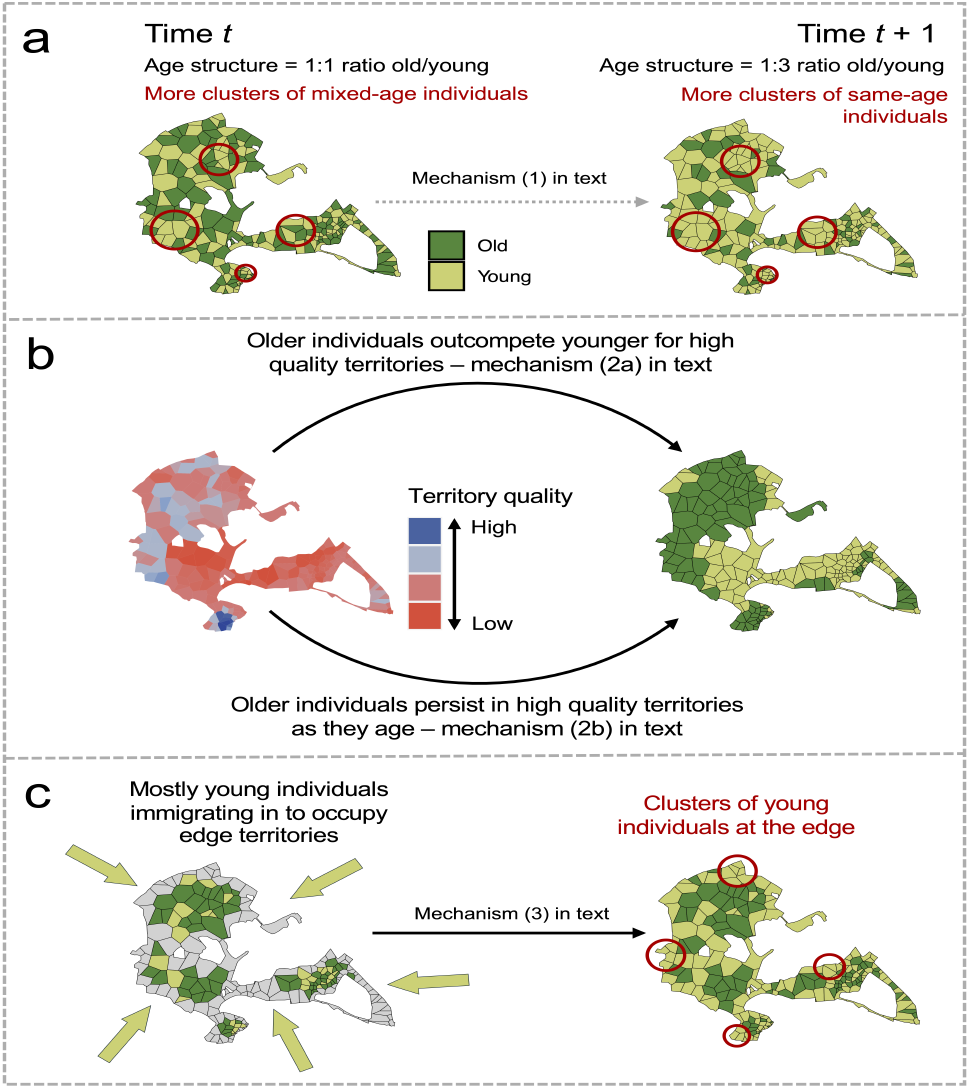
Overview of mechanisms that might generate spatial age structure across territories referred to in the introduction. The theoretical populations consist of 200 individuals, each of which are associated with a territory, illustrated across Wytham Woods, Oxford, as an example. Yellow and green territories are occupied by ‘young’ and ‘old’ individuals respectively. In (a), individuals occupy territories randomly, but at time *t* the population consists of a 1:1 ratio of old/young individuals; and at time *t + 1* a 1:3 ratio of old/young individuals. In (b), the population consists of a 1:1 ratio of old/young individuals, but older individuals occupy the territories of highest quality. In (c), territories which do not border the edge are occupied by a 3:1 ratio of old/young individuals, whereas territories with an edge boundary are occupied by a ratio 1:3 of old/young individuals.

However, age-specific biases in settlement may drive spatial age structure over-and-above that expected from temporal variation in age distribution alone. This may occur if there is spatial structure in territory quality (which may be generated for example through spatial clustering of resources), and there is covariance between age and territory quality, which may arise through two mechanisms. (2a) In many species, older individuals are dominant (Piper 1997; Wilson 2000), thus potentially leading to age-specificity in territory use where older individuals outcompete younger ones to acquire higher quality territories (Ferrer and Bisson 2003; Garneau et al. 2008; Brewer et al. 2009; Stepanuk et al. 2021) (Fig 1b). (2b) Alternatively, covariance between age and territory quality might arise if higher quality territories elevate survival, and individuals therefore persist for longer in such territories over time (Kokko et al. 2006). Additionally, individuals might be more likely to remain on higher quality territories, which will generate a similar association between length of tenure and hence age (Fig. 1b). Therefore, if older individuals are more likely to occupy higher quality territories through either of these mechanisms, spatial structure in territory quality might drive spatial age structure and repeatability in this over time.

(3) Age-specific territory occupation might also be generated through mechanisms that do not directly link to how ageing affects territory acquisition. For example, if there is individual-specificity in territory acquisition mediated by states or phenotypes other than age, but there is covariance between these and age, then association may arise between age and the type of territory (despite no causal direct link between the two). For example, territory-specific occupation depending on prior dispersal is relatively common across taxa (Piper 2011). Specifically, in fragmented landscapes, sites have environmental ‘edges’, such as forest edges which provide an interface between internal woodland and the external environment (Saunders et al. 1991). In edge environments, there is often a higher incidence of individuals that disperse into the site (Krauss et al. 2003; Wilkin et al. 2007a). Dispersing between birth and first breeding, known as natal dispersal, generally involves greater distances than dispersal between two breeding attempts (Greenwood and Harvey 1982; Johst and Brandl 1999). Thus, individuals moving into a new environment are often younger (Greenwood 1980; Verhulst et al. 1997; Slagsvold and Wiebe 2018). In such cases, if edge territories are more commonly occupied by younger immigrant individuals, then this might provide a mechanism through which spatial age structure develops (Fig. 1c). Edge territories provide a particularly interesting mechanism by which spatial age structure is generated, not only because of immigrant-specific occupation, but also because they are often of poorer quality (Wilkin et al. 2007a; Purcell et al. 2012; Loveridge et al. 2017), thus they might also directly generate spatial age structure if there is covariance between age and territory quality (mechanisms 2a and 2b).

These mechanisms lead to *a priori* expectations that spatial age structure might develop across breeding territories; however, empirical evidence of this is currently sparse. Additionally, although these mechanisms might act concurrently on individuals in time, the areas in space that bias age-specific settlement may differ from each other. For example, if different aspects of territory quality are independent then the relative strength of these resource distributions, if any, in influencing age-specific settlement is unknown. Further, how such mechanisms bias age-specific settlement might be influenced by sex given the different roles that males and females will often play in territory acquisition (Brown 1964; Greenwood 1980). In short, given the role that age structure might play in spatially-structured demography, it is important that we advance our understanding on the evidence for spatial age structure and the mechanisms that drive it.

Here, we outline processes and their consequences for generating spatial age structure on long-term breeding data from a natural population of great tits Parus major in Wytham Woods, Oxford. We first assess whether there is covariance between territory quality and the age of individuals in such territories. Second, we evaluate whether there is non-random spatial arrangement of age within breeding seasons. Finally, we assess evidence for age-biases in space that persist across time compared to spatial structure in territory quality and reproductive output. We discuss the relative roles of spatial age structure versus structure in territory quality for producing spatially-variable demography, and suggest that the framework presented here can be used across populations when considering spatial age structure.

## 2 Methods

### 2.1 Study system and data collection

The great tit *Parus major* is a passerine bird found in woodlands across Europe, with breeding ages ranging 1–9, averaging 1.8 years (Perrins 1979; Gosler 1993; Bouwhuis et al. 2009). Although there are some continuous changes with age (Bouwhuis et al. 2009), the main age effects on individual-level traits are captured by two age-classes: first-years (hereafter juveniles) and older (hereafter adults, Greenwood et al. 1979; Perrins 1979; Sandell and Smith 1991; Gosler 1993; Farine et al. 2015). The species is socially monogamous, with pairs defending territories during annual breeding seasons (Hinde 1952). Data used here are from a long-term study in Wytham Woods, Oxford (51°46’N, 1°20’W), a 385ha mixed deciduous woodland surrounded by farmland (Savill et al. 2010). The tit population has been monitored since 1947, where breeding adults and their chicks have been marked with unique BTO (British Trust for Ornithology) rings since the 1960s; and standard reproductive metrics are collected (Perrins 1965). Individuals breed almost exclusively in the 1026 nest-boxes which are in fixed positions with known GPS coordinates (Krebs 1971; Wilkin et al. 2006). All chicks are ringed at 14-days of age, while adults are trapped at nest-boxes and identified by ring number, or marked with a new ring if they are immigrants. Age is based on year of hatching for local birds, or plumage characteristics for immigrants (Svensson 1992). Although immigration rates are high (46%), most are first caught as yearlings (78%) and can therefore be aged accurately.

### 2.2 Data selection

We constructed a dataset that assigns the year of hatching to all individuals between 1950–2022, across which exact age was calculated for 88.8% of 46062 identified breeding individuals. In this study, we included birds in analyses that attempted to breed between 1978–2022, for which data were more complete compared to earlier dates (Fig. S1). Individuals that were first caught post-fledging are assumed to be immigrants, as locally-hatched tits are marked as nestlings in nest-boxes and the proportion of birds hatched in natural cavities is very low (Kidd et al. 2015). Immigrants that entered the population with adult plumage were assigned a minimum age of 2, and subsequent age estimates were based on this (6.7% and 10.0% of breeding females and males). Age was therefore determined for 68.7% of breeding individuals where at least one egg was laid (due to a combination of nests failing prior to adult trapping and unsuccessful trapping attempts, there are cases where the identity of parents is unknown).

### 2.3 Statistical analyses

#### 2.3.1 Determining breeding territories

We defined annual breeding territories through a Dirichlet tessellation technique that forms Thiessen polygons (Rhynsburger 1973; Tanemura and Hasegawa 1980) around each occupied nest-box. The polygon includes all space within the habitat that is closer to the focal box than any other (with a boundary also imposed by the woodland edge). This metric of territory has been shown to be biologically meaningful in terms of territory size and territorial neighbours in tit species and is strongly related to other methods of calculating territories (Adams 2001; Wilkin et al. 2006; Schlicht et al. 2014; Firth and Sheldon 2016; Gokcekus et al. 2023). However, a limitation is that unrealistically large polygons are formed in areas where nest-boxes are placed at great distances from each other. We therefore capped territories at 2ha, which is a more realistic maximum spatial scale at which individuals use territories, as supported in previous analytical and field studies (Krebs 1971; Both and Visser 2000; Wilkin et al. 2006, 2007a).

#### 2.3.2 Age and territory quality

We first assessed covariance between the age of individuals and their territories’ quality. We measured territory quality through four measures: the number of oak trees *Quercus* spp. within 75m of the nest-box; average territory density; the edge distance index; and the long-term nest-box popularity index. Each of these is justified below.

Great tits predominantly provision offspring with caterpillars collected close to their nests (Gosler 1993), thus variation in caterpillar availability is directly linked to reproductive success (Perrins 1991; Keller and van Noordwijk 1994; Rytkö nen and Krams 2003). Caterpillars are found most abundantly on oak trees (Perrins 1991; Gosler 1993), therefore oak proximity, health and abundance is important for breeding success (Wilkin et al. 2006, 2009; Cole et al. 2021; Gokcekus et al. 2023). A radius of 75m was chosen as the abundance of oaks within this distance has been shown to be particularly important for breeding (Wilkin et al. 2007b; Hinks et al. 2015).

The density of conspecifics breeding in proximity may influence resource availability if foraging ranges overlap, and therefore territory density may also represent an aspect of territory quality. Additionally, territory density may affect site quality through social mechanisms, such as increased competition and emergent need for territory defence leading to reduced foraging (Ydenberg and Krebs 1987), or conversely mutual benefits between familiar neighbours (Grabowska-Zhang et al. 2012b, 2012a; Gokcekus et al. 2023). We calculated average territory density directly from the Thiessen polygon area produced from tessellation by taking the reciprocal of the mean polygon area.

Territories at woodland edges are associated with lower reproductive success in great tits (Wilkin et al. 2007a). Following Wilkin et al. 2007a, we defined the edge distance index (EDI) for each nest-box by multiplying the distance to forest edge by the proportion of woodland habitat within a 75m radius of the box. Thus, boxes within 75m of the edge have an EDI value in proportion to the amount of woodland habitat within this radius, therefore considering not only the distance to edge, but also the number and geometric arrangement of edges relative to nest-box.

Finally, the frequency a territory is occupied in the long-term may provide a measure of quality as individuals may choose sites that confer reproductive benefits more often, as evidenced in other species (Sergio and Newton 2003; Smith and Moore 2005; Johnson 2007; Steenhof and Newton 2007; Potti et al. 2018). There is evidence of this in Wytham, where the number of times a nest-box has been occupied positively correlates with the average number of offspring that fledge per breeding attempt (Fig. S2). We therefore calculated the number of times a nest-box has been occupied since 1965. However, this is related to the number of nest-boxes within close proximity to a focal box (Fig. S3), because in areas of high nest-box density there are multiple unoccupied boxes which would likely be associated with the same territorial range if they were occupied (thus, in regions of high box density, birds may re-occupy the same territory over multiple years, but not necessarily the exact same box). To correct for this, we ran a linear model between the number of boxes within 30m of a focal nest-box (supporting information) and the number of times said box has been occupied, and took the residuals as the long-term nest-box popularity index.

We constructed a generalised linear mixed-effects model assuming a binomial error distribution to analyse the association between these four measures of territory quality and the age of the breeding individual. We modelled age (juvenile/adult) as the response variable, with the territory quality measures as explanatory variables, which were z-transformed to compare their relative effects in predicting age. Individual ID, nest-box ID, and breeding year were included as random effects (Table S1). We ran three sets of these models: one with all individuals; one with only females; and one with only males, allowing us to assess potential sex-specific differences in the association between aspects of territory quality and age. These models were also run to test for covariance between territory quality and residency status (supporting information). All models were conducted in R statistical software (R Core Team 2021) using the *lme4* package (Bates et al. 2015).

#### 2.3.3 Spatial age structure

For each year’s breeding population, we constructed an individual-by-individual matrix denoting breeding neighbours i.e. a network of breeding territories, where nodes represent individuals and edges represent the spatial connectivity of territories. Specifically, edges connected individuals if their territories share a boundary from the tessellation technique and were weighted relative to the distance between nest-boxes of neighbouring territories.

We created three networks per year: one with all individuals (but removing edges within breeding pairs that occupy the same territory); one with only females; and one with only males. Edges connecting individuals of unknown identity were removed. Across these networks, we calculated the assortativity coefficient of age (juvenile/adult), which measures the correlation between individuals’ age and that of their territorial neighbours accounting for edge weight (proximity of neighbouring nest-boxes) and the relative proportion of the two age classes across the network. Through this technique, the age-composition of neighbourhoods of birds that are likely to interact during territorial and foraging behaviour contribute to the emergent quantitative signal of spatial age structure to a greater degree than they would if spatial autocorrelation were calculated through pairwise distance of all individuals across the system. Thus, this method allows us to assess evidence for spatial age structure at a biologically relevant scale (analyses were also run treating age as a discrete and continuous trait, as well as calculating assortment of residency status; supporting information). We also ran analyses to compare spatial age structure with spatial structuring of territory quality and reproductive output. To calculate territory quality structure, we assigned the value from the previously described four measures of quality to each node (associated with an occupied nest-box) and calculated the territory quality assortativity coefficient across the network. For spatial reproductive output structure, we ran parallel analyses, but calculated the assortativity in clutch size, chick number, fledgling number, and binary success (where 0 is no fledglings and 1 is at least one fledgling) associated with each nest-box. These analyses were run using the *assortnet* package (Farine 2014).

#### 2.3.4 Temporal repeatability of age structure

Finally, we tested whether spatial regions show temporal repeatability in age-composition, territory quality, and reproductive output. To assess this, within each year, we defined a radius around each occupied nest-box that corresponded to an area of 25, 50, 100, 150 and 200ha, representing neighbourhoods of breeding individuals of variable population sizes. Within each radius we calculated: the proportion of individuals that were adults; the mean number of oaks within 75m of the focal boxes; mean territory density; mean edge distance index; mean nest-box popularity index; mean clutch size; mean chick number; mean fledgling number; and proportion of boxes with binary success. We then calculated the same metrics for all nest-boxes outside of the focal radius. From this, we calculated a ratio index of within versus outside radius for each calculated measure, where a value of one represents the same annual average measure within and outside the radius. We then tested the repeatability of the ratio index associated with each spatial scale across the 45 years of data by calculating the intraclass correlation coefficient (ICC) derived from a linear mixed-effects model, where the response variable is the ratio index, and year and radius area are fitted as categorical grouping variables.

## 3 Results

### 3.1 Age and territory quality

Across 20121 breeding individuals, representing 11167 breeding attempts, we found that high density territories weakly predicted occupation by adults compared to juveniles (odds ratio = 1.108, 95% confidence intervals = 1.065–1.153, p *<* 0.001), in addition to territories with a greater long-term popularity index (OR = 1.084, 95% CIs = 1.045–1.124, p *<* 0.001), with little evidence that the number of oaks within 75m or EDI were linked to age (Fig. 2; Table S2). When assessing the influence of territory quality on the age of individuals separately for both sexes, similar patterns were found, except territories that have a greater long-term popularity index more strongly predicted occupation by adult males (OR = 1.143, 95% CIs = 1.092–1.195, p *<* 0.001), but were weakly if at all associated with female age (OR = 1.041, 95% CIs = 0.997–1.086, p = 0.066).

**Fig. 2.**
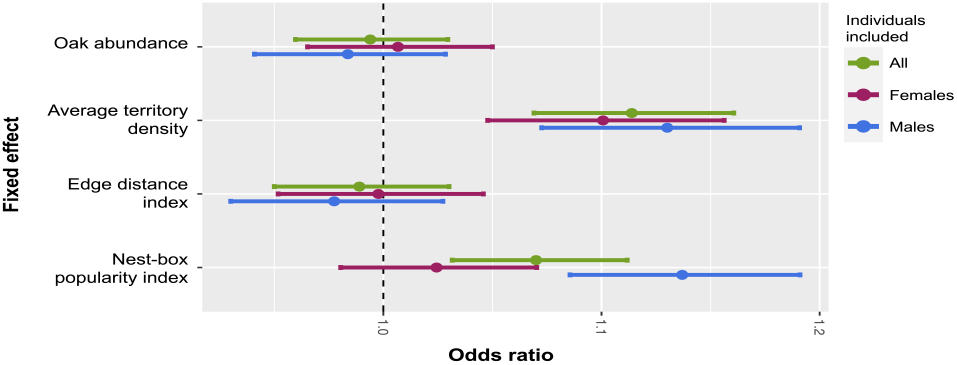
Point estimates of odds ratios obtained from generalised linear mixed-effects models. The analysis examines the association between territory quality on the odds of the breeding individual being an adult. Each point corresponds to a specific level of the fixed effects (on the y-axis), and error bars denote 95% confidence intervals. Green points are from analysis assessing all individuals, purple are only females, and blue are only males.

### 3.2 Spatial age structure

Generally, we found weak positive age assortment across breeding territories, but there was temporal variation with many years having little evidence for spatial age structure and some having significant disassortment (i.e. birds of similar age less likely to be associated), whether considering the whole population (median r age assortativity coefficient = 0.014, 51% of annual standard errors overlap zero; Fig. 3), females (median r = 0.002; 58% SEs overlap zero), or males (median r = -0.004; 38% SEs overlap zero). In contrast, there was much greater spatial structure in territory quality (oak abundance median r = 0.782; territory density median r = 0.645; EDI median r = 0.817; and nest-box popularity index median r = 0.258, no annual SEs overlap zero). There is greater evidence for spatial reproductive output structure compared to age, but the signal of this structure is still relatively weak (clutch size median r = 0.059, 22% SEs overlap zero; chick number median r = 0.047, 29% SEs overlap zero; fledgling number median r = 0.075, 22% SEs overlap zero; binary success median r = 0.057, 24% SEs overlap zero; Fig. 3; Table S3 for all results).

**Fig. 3.**
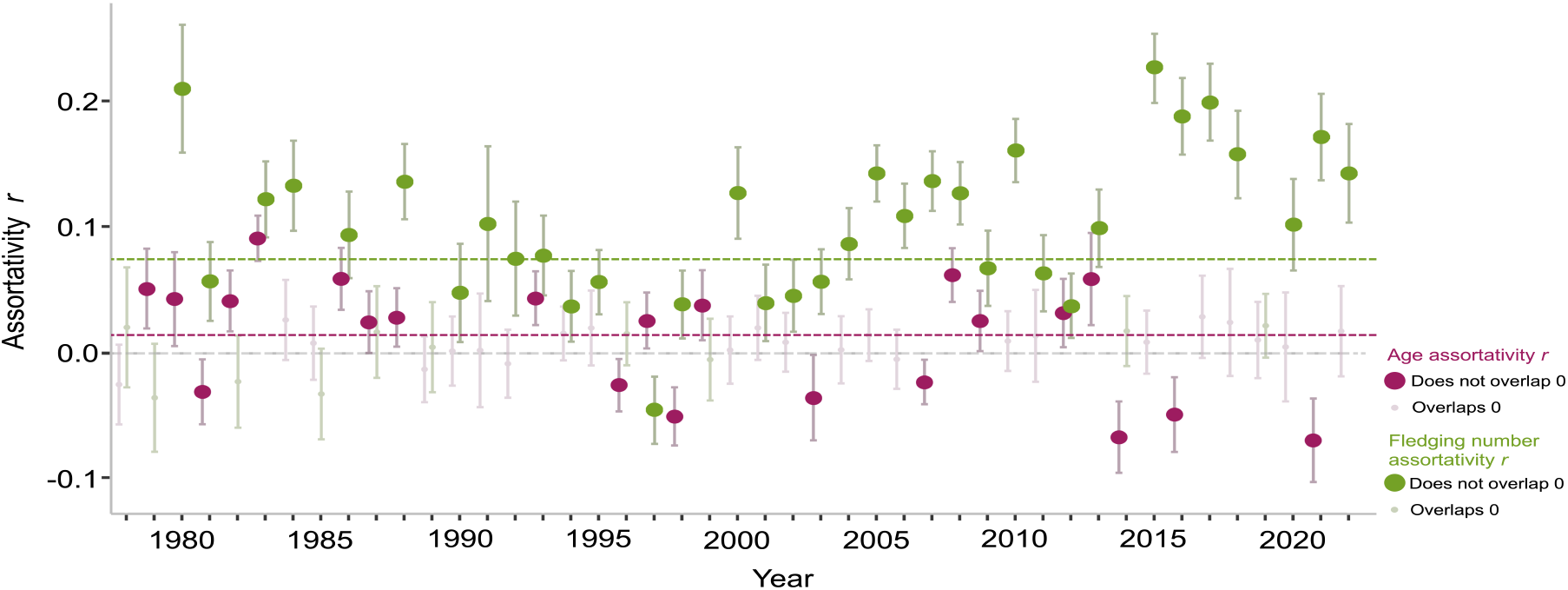
Annual age and fledgling number assortativity r coefficients across territories. Large purple points represent age assortativity values where the standard error does not overlap zero, and large green points are fledgling number assortativity values where standard error does not overlap zero. The dotted purple line denotes the median age assortativity value, and the green line denotes the median fledgling number assortativity value.

### 3.3 Temporal repeatability of spatial age structure

There was low temporal repeatability in the age-composition of spatial regions at all assessed scales 25–200ha (ICC range: 0.036–0.057; Fig. 4). In contrast, as would be expected, there is very high temporal repeatability in all assessed territory quality measures (Fig. 4). Compared to age-composition, there is higher temporal repeatability in average reproductive measures of neighbourhoods, with mean clutch size (ICC range: 0.234–0.447) displaying the highest repeatability (Fig. 4; Table S4 for all results).

**Fig. 4.**
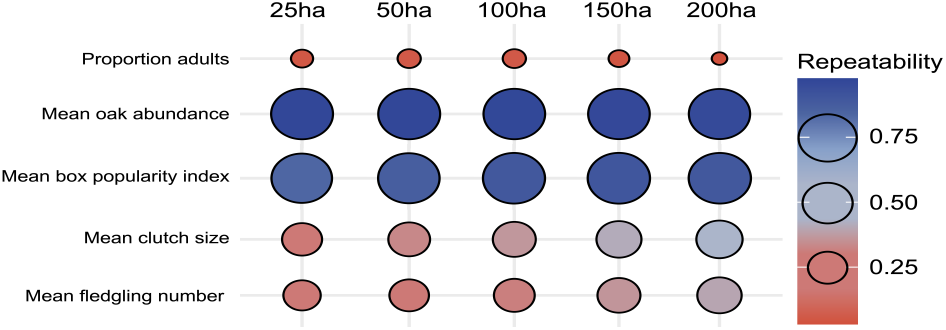
Temporal repeatability of age-composition, territory quality and reproductive output of neighbourhoods at spatial scales of 25, 50, 100, 150 and 200ha. Repeatability was derived from the ICC from a linear mixed-effects model, where the ratio index of within versus outside the defined spatial scale of the average measure was the response variable, and year and spatial scale area categorical grouping variables. Circles are sized and coloured on a gradient from red to blue as repeatability increases.

## 4 Discussion

We quantified the extent of spatial age structure in a large long-term study population of territorial birds, and tested several mechanisms that could generate assortment by age in space. We show that, in general, spatial age structure is weak, despite the occurrence of spatial structure in habitat features, some of which covary with age and could plausibly generate spatial age structure. We also find that there is little temporal repeatability of spatial regions which bias certain age structures, despite stronger evidence for repeatability in spatial structure of reproductive output and territory quality. Here, we discuss these findings, suggesting that spatial structure in territory quality might be a greater contributor to spatially-variable demography and why there is not greater spatial age structure in this system.

### 4.1 Age and territory quality

Across all breeding individuals, we show that older individuals are more likely to acquire territories at higher densities that have a greater long-term popularity, while there is a lack of evidence for a relationship between age and the abundance of oak trees or distance to edge. Considering females and males separately, we find similar associations between age and territory quality, except that the popularity index of a nest-box more strongly predicts occupation by adult males than females.

It is interesting that territories at higher average densities are more likely to be occupied by adults than juveniles considering that this may result in reduced territory size, increased chance of encounter with conspecifics during chick-rearing behaviour, and has previously been linked to lower reproductive success (Wilkin et al. 2006). This may relate to evidence that adults have a weaker response to density dependence in great tit populations, where juvenile cohorts have a greater effect in reducing recruitment and survival, and more negatively respond to density dependence (Gamelon et al. 2016). Together with our results, this may reflect improved breeding competence of adults compared to juveniles leading to higher territorial densities. Specifically, older individuals may have increased foraging efficiency (Perrins and McCleery 1985), thus negating the need for large territories (indeed, individuals with smaller foraging ranges can be more efficient with higher reproductive success, Cole et al. 2012), therefore generating higher breeding densities among adults and a weaker response to density dependence. Additionally, increased familiarity between adult territorial neighbours may reduce competition associated with foraging (Grabowska-Zhang et al. 2012b; Gokcekus et al. 2023), or even promote joint benefits such as nest defence (Grabowska-Zhang et al. 2012a), thus favouring closer breeding between adults. Recent advances in tracking technology of passerine foraging behaviour (Baldan and van Loon 2022) might advance insight into whether competition is reduced among older familiar neighbours by assessing evidence for increased coordination during foraging compared to unfamiliar neighbours.

Territories that were more popular in the long-term were more likely to be occupied by adults than juveniles. The use of frequency of occupancy as a measure of territory quality is well-established in avian systems (Sergio and Newton 2003; Smith and Moore 2005; Johnson 2007; Steenhof and Newton 2007; Potti et al. 2018), and may capture information on cues of quality which individuals respond to in the long-term which cannot be directly measured. It is particularly interesting that the association between more popular territories and age is weakened when only assessing females, but strengthened in males. This may be explained by sex differences where male great tits are more important in initial acquisition and defence of territories (Krebs 1971, 1977; Perrins 1979; Bjö rklund et al. 1989), which is widespread in other avian (Brown 1964; Stutchbury 1991; Beletsky 1992; Eikenaar et al. 2009) and taxonomic groups (Clutton-Brock 1989; López and Martín 2002; Milner et al. 2010). Thus, this could suggest that older males with greater competitive ability (Sandell and Smith 1991) outcompete younger individuals to occupy higher quality sites. This may be driven either by novel acquisition of high-quality territories, or by persistence and defence of a territory between years (Greenwood et al. 1979; Harvey et al. 1979a). In the latter case, males may occupy a high-quality territory in their first breeding attempt, which supports successful breeding, and then retain this territory in future breeding. Thus, if survival is high between years, particularly among individuals in high-quality territories, then covariance between male age and territory quality may arise as younger first-year breeders are forced into low-quality territories.

There is little evidence that the abundance of oaks close to the nest predicts age. Oak abundance at this scale has been linked to great tit breeding success (Wilkin et al. 2006, 2007b, 2009; Hinks et al. 2015; Cole et al. 2021; Gokcekus et al. 2023), thus increased experience and competitive ability of adults (Sandell and Smith 1991) might lead to the assumption that older individuals will acquire territories with more oaks. However, although higher oak abundance might provide better quality territories when considering entire populations, little is known about how this interacts with phenological synchrony at the individual-level (Hinks et al. 2015) and how the degree of phenological matching depends on age. Specifically, older great tits typically breed earlier (Harvey et al. 1979b; McCleery and Perrins 1998), possibly due to improved breeding competence leading to greater synchrony between chick-rearing and peak caterpillar abundance. Further, the rate at which caterpillars are provisioned to chicks strongly correlates with the caterpillar biomass available (Naef-Daenzer et al. 2000). Thus, if a greater proportion of juveniles mistime their breeding compared to adults, then they may be unable to monopolise caterpillar resources even if in oak-rich territories due to temporal mismatch. In addition to greater phenological synchrony, older birds may improve their foraging ability in a broader sense through greater site familiarity from previous occupation (Perrins and McCleery 1985; Piper 2011; Slagsvold and Wiebe 2018), utilisation of a wider variety of food resources (Gosler 1993), and improved foraging coordination with pair-bonded partners (Mariette and Griffith 2015; Griffith 2019). In this sense, the overall abundance of oaks may not be a crucial cue when acquiring territories, rather there is stronger selection on phenological matching to the local environment (Hinks et al. 2015; Cole et al. 2021) and general foraging ability. Additionally, there is little evidence that the edge distance index is associated with age, which has been linked to reduced reproductive output (Wilkin et al. 2007a), and thus there is no apparent age-specificity in how edge effects mediate territory acquisition.

The lack of an effect of oak abundance and edge distance on age-specific territory occupation, as well as the general weak association with other territory attributes, may suggest that territory acquisition in adults is predominantly driven by retaining previous territories in which they successfully bred, regardless of quality. This may be driven by a relative lack of available territories compared to overall population size, thus providing strong selection for the retention of a previously occupied territory (Clark 1994; Harts et al. 2016). High breeding site fidelity is evident in great tits (Harvey et al. 1979a). In this case, the cue which individuals respond to when acquiring a territory may be experience of previous breeding success, as opposed to current environmental quality cues. Consequently, any covariance between older age and high territory quality might break down through two mechanisms: either previous breeding occurred in a low-quality territory, but breeding was successful and thus the territory is retained as an adult; or previous breeding occurred in a high-quality territory, but the attempt was unsuccessful, and thus individuals move to new poorer quality territories in future years. Breeding success in poor quality territories can regularly occur, for example if intraspecific competition is low, or when individual or partner quality is high. In such cases, older individuals might utilise their higher competitive ability to defend territories in which they have had success regardless of quality (Harvey et al. 1979a; Yasukawa 1979; de Vos 1983). Conversely, unsuccessful breeding may happen in areas of high-quality, through stochastic environmental effects (such as extreme weather or predation), or low individual or partner quality. In these cases, despite being in high-quality territories, pairs may divorce in an attempt to improve future breeding (Culina et al. 2015) and disperse to new territories (Harvey et al. 1979a). Through this process, older individuals will not necessarily occupy the best quality territories, thus leaving unoccupied territories of higher quality for juveniles to acquire.

Given our findings, future research should aim to disentangle the relative effects of territory quality and previous reproductive success on site fidelity in territorial species, as well as assessing behavioural differences in the degree of territory defence in individuals that re-occupy sites of varying quality. Further, examining age-specific behaviour following divorce and acquisition of a new territory might shed light on how age relates to territory quality when removing the effect of site fidelity. For instance, there may be variation in territory quality across divorced individuals that newly settle depending on their own age and that of their new partner. Finally, more work should assess age-specificity in territory acquisition while considering non-breeding individuals that fail to acquire a territory. For example, in great tit populations, there are often more individuals than available territories (Perrins 1979), leaving non-breeders in the population during the spring. In avian species, non-breeders tend to be younger (Stutchbury 1991; Sergio et al. 2009) and the ability to defend a territory against intrusions from non-breeders increases with age (Arcese 1987). Thus, experiments to identify non-breeders during territorial activity (Firth et al. 2018) and accounting for this cohort might reveal further age-specificity in territory quality.

### 4.2 Spatial age structure

Our study reveals more frequent positive annual spatial age structure, but this structure is weak and often absent, with no clear sex differences. In contrast, there is strong evidence for spatial territory quality structure, with particularly strong assortment in oak abundance, territory density and edge distance (the latter of which is spatially necessitated), and moderate assortment in the long-term popularity index of a nest-box. There is greater evidence for spatial structure in reproductive output compared to age, with fledgling number displaying the greatest structure.

It is interesting that there is not greater spatial age structure not only due to our evidence of covariance between some attributes of territory quality and age, and spatial structure in such territory quality, but also due to social mechanisms that might generate spatial age structure. Winter social associations in great tits are positively age-assorted, albeit fairly weakly (Farine et al. 2015), and social connections carryover into breeding as individuals establish territories closer to previous social associates (Firth and Sheldon 2016). This is likely because having familiar neighbours can reduce the need for territory defence, increase cooperation, and enhance reproductive success (Grabowska-Zhang et al. 2012b, 2012a; Gokcekus et al. 2023). Thus, because older individuals are more likely to be familiar with their neighbours than younger ones (Gokcekus et al. 2023), this might be expected to generate spatial age structure through clusters of older individuals breeding in proximity. However, although this mechanism provides direct benefits at the dyadic-level for closer breeding between adults, there is little overall spatial age structure. This may be partly explained by the species’ life-history, where there is high annual mortality (52%, Bouwhuis et al. 2009). Together with annual influxes of immigrant birds, this means that the breeding population is often composed of a large proportion of individuals that are first-time breeders (46–83%, Woodman et al. 2022), thus only making it possible for a small proportion of the population to have familiar neighbours. In short, despite benefits for older breeders to retain previous neighbours, high mortality rates lead to a high chance that at least some previous territorial associates will have died, and high competition for breeding sites mean these will be acquired by new and likely juvenile breeders (Krebs 1977; Perrins 1979; Bjö rklund et al. 1989), thus reducing overall spatial age structure. This hypothesis should be tested through comparative analyses using the framework developed here by assessing whether there is greater spatial age structure in territorial species with lower mortality rates and thus a greater opportunity to retain previous neighbours, particularly considering familiarity between neighbours is adaptive in many taxonomic groups (Temeles 1994; Frostman and Sherman 2004; Siracusa et al. 2021).

Additionally, although there are benefits of retaining familiar neighbours, this is not necessarily underpinned by an active process where older individuals select territories based on closer breeding to previous associates. Instead, it may be underpinned by a passive process where individuals which survive between years occupy the same locality (Harvey et al. 1979a), leading to the incidental retention of familiar neighbours that also survive and have high site fidelity. Such a passive mechanism is consistent with recent findings that age-assortative mating within pairs of great tits, despite being adaptive (Harvey et al. 1979b), is underpinned by passive processes of pair retention and fluctuations in overall population age structure as opposed to active age-related selection (Woodman et al. 2022). Thus, even if there are direct advantages of retaining familiar neighbours, if it is not underpinned by active processes, then this might weaken its prevalence and therefore overall spatial age structure.

Finally, although there is temporal overlap in when territories are used, there is variation between pairs which will affect how neighbours interact depending on their breeding time synchrony, thus affecting spatial age structure at any single point in time. Firstly, there is spatial variation in the timing of food availability at a spatial scale relevant to individual territories, underpinned by variation in individual tree phenology (Cole and Sheldon 2017), and this local variation predicts breeding timing in individual pairs (Hinks et al. 2015). Thus, pairs with neighbouring territories may not interact much during chick-rearing behaviour if neighbouring phenology is highly asynchronous (although there is generally spatial structure in breeding phenology across Wytham, Wilkin et al. 2007b). Second, there might be even greater temporal mismatch in occupation of neighbouring territories when considering initial acquisition and defence behaviour, which begins in January or even earlier, months before chick-rearing (Krebs 1977, 1978, 1982). Thus, interactions between individuals and competition associated with territory defence will be variable depending on the occupation status of neighbouring territories at the time of focal territory acquisition. For example, there may be extended periods of territorial activity pre-breeding where neighbouring pairs interact, but then a new pair may acquire a territory which bisects such breeding sites, thus breaking down the territorial boundary between the two original neighbouring pairs. Previous research in this system has assessed age-specificity in nest-site visitation patterns prior to breeding in terms of the number and spatial clustering of nest-boxes visited (Firth et al. 2018). Thus, future work could incorporate such techniques with advances in the tracking of movement (Levin et al. 2015; Baldan and van Loon 2022) and other defence behaviour (Merino Recalde 2023) to advance our understanding on the temporal scale at which individuals interact during territorial behaviour before breeding, how this is affected as territories are gradually occupied prior to the spring, and age-specific variation in this.

### 4.3 Temporal repeatability of spatial age structure

We find low repeatability in the age-composition of spatial regions between years compared to repeatability in spatial territory quality and reproductive output structure. High repeatability in spatial territory quality structure is expected, as annual variation in average quality measures across spatial regions is changed only depending on which nest-boxes are occupied (i.e. if the same boxes were occupied every year, there would be 100% repeatability). However, there is variation in which boxes are used each year. Thus, in conjunction with our previous finding of high within-year spatial territory quality structure, it is interesting then that we report a higher repeatability in the average reproductive output compared to the age-composition of spatial regions. This suggests that territory quality in this system may be more important in driving spatially-variable demography than local age structure. Additionally, the finding of low repeatability in spatial age-composition is consistent with our suggestion that adult occupation is likely largely driven by persistence in previous territories of varying quality as opposed to active acquisition of high-quality territories (the latter would lead to higher repeatability as older birds would frequently use higher quality sites independent of year).

## 5 Conclusions

Our study highlights association between age and aspects of territory quality in a territorial passerine species, specifically showing that older individuals breed at higher densities and in sites which are more popular in the long-term. However, we show limited evidence of spatial age structure. We suggest that age-specific occupation of territories is likely driven by persistence in territories across lifespan (largely irrespective of quality), leading to passive retention of some older neighbours which also survive between years. We emphasise the importance of exploring the operation of these mechanisms that might lead to spatial age structure in species with different life-histories. Specifically, it might be expected that in species with lower mortality rates, greater spatial age structure would be generated through passive mechanisms of territorial neighbour retention as more individuals survive between years. The observed patterns of higher spatial territory quality and reproductive output structure (compared to spatial age structure), in addition to greater temporal repeatability in these, contributes valuable insights into the drivers of spatially-variable demography. Namely, given these results, we suggest that spatial clustering of territory quality is likely to be more important in determining repeatable spatial structure in reproductive output compared to spatial age structure.

## Acknowledgments

We thank the many Wytham Woods fieldworkers who have worked on the tit study for the past 76 years.

