## Supplementary material for "Age-specificity in territory quality and spatial structure in a wild bird population": Figure S; supporting information; Table S

4 Supplementary figures 4

5 Supplementary tables 8

6 References 27

#### 1 SUPPLEMENTARY METHODS

##### 1.1 Nest-box density and occupation

The number of times a nest-box has been occupied by a great tit since 1965 is likely affected by the density of nest-boxes placed in close proximity. This is because during the breeding season, in areas of high box density, there are multiple unoccupied boxes which would likely be associated with the same territorial range if they were occupied (Fig. S4), which is variable between individuals but estimated as approximately 2ha in area or within a 75m radius around a focal box (Krebs 1971; Barnes 1975; Both and Visser 2000; Wilkin et al. 2006, 2007b, 2007a; Hinks et al. 2015). Therefore, in areas of high nest-box density, birds may re-occupy effectively the same territory over multiple years, but not necessarily the exact same box, which is not the case in areas of low box density. To correct for this when using occupation rate as a measure of territory quality, we ran a linear model between the number of boxes within 30m of a focal nest-box and the number of times said box has been occupied, and took the residuals as the corrected long-term nest-box popularity index. 30m was chosen as territorial ranges (defined by a radius of 75m) around two focal points placed at 30m or less from each other overlap by 75% or more (Fig. S4), thus largely representing the same territory. While it is currently unknown at which precise distance between neighbouring boxes an inter-annual switch represents retaining largely the same territory for breeding tits (as well as the fact that this is likely to be variable depending on individual behaviour and local population size), previous analytical and field studies suggest that 30m is an appropriate distance to use based on our previous justification of territorial ranges. However, we additionally defined the nest-box popularity index by taking the residuals of a linear model between the number of boxes within 10, 20, 40, 50 and 75m of a focal nest-box and the number of times occupied. This did not substantially change the point estimates of our results when running a generalised linear mixed-effects model to test the association between our measures of territory quality and the age of the individual in a territory (Table S3).

##### 1.2 Methods assessing residency status

###### *1.2.1 Residency status and territory quality*

Due to the association between residency status and age where a high proportion of immigrants are juveniles, in addition to well-established biological differences between residents and immigrants where residents are socially dominant (Sandell and Smith 1991; Farine and Sheldon 2015; Slagsvold and Wiebe 2018), have higher reproductive success (Dhondt et al. 1990; Wilkin et al. 2007a; Slagsvold and Wiebe 2018) and have greater site familiarity (Krebs 1982; Wilkin et al. 2007a; Slagsvold and Wiebe 2018), we ran parallel analyses to assess whether territory quality measures predict residency status. We define two types of residency status: immigration status (immigrant/local recruit: an individual-level trait that remains the same throughout life-history depending on whether the bird hatched within or outside of Wytham); and settled status (new/settled; an individual-level trait which may change across lifespan, where new birds refer to immigrants that are breeding for the first time, Fig. S5).

We therefore constructed a generalised linear mixed model assuming a binomial error distribution as in the main text to analyse the association between the four measures of territory quality and the immigration/settled status of the individual that acquires and breeds in a territory. Three models were run for each of these two binary traits: one with all individuals; one with only females; and one with only males.

###### *1.2.2 Spatial structure in residency status*

As in the main text, we assessed spatial structure in residency status by creating three networks per year: one with all individuals (but removing edges between breeding pairs that occupy the same territory); one with only females; and one with only males. Across these networks, we calculated the assortativity coefficient of immigration status (immigrant/local recruit) and settled status (new/settled).

###### *1.2.3 Temporal repeatability of spatial structure in residency status*

We tested the scale at which regions in space show repeatability in spatial structure in immigration and settled status. For each year, within each radius associated with a focal nest-box as outlined in the main text we calculated the proportion of breeding individuals that are local recruits and the proportion that are settled birds, and also calculated this for all nest-boxes outside the focal radius. We then tested the repeatability of the ratio index associated with each variably sized radius across the 45 years of data collection by calculating the intraclass correlation coefficient (ICC) derived from a mixed-effect model, where the response variable was the ratio index, and year and radius area were fitted as random categorical grouping variables.

#### 2 Supplementary Results

##### 2.1 Territory quality and spatial structure in residency status

###### *2.1.1 Residency status and territory quality*

Territories at lower densities significantly predicted occupation by local recruits compared to immigrants across all individuals (OR = 0.662, 95% CIs = 0.567–0.773, $p < 0.001$) and in females (OR = 0.624, 95% CIs = 0.504–0.773, $p < 0.001$), but not in males (OR = 0.951, 95% CIs = 0.748–1.210, p = 0.683; Fig. S6; Table S2). There is little evidence that that the long-term popularity index or the abundance of oaks predicted the immigration status of the breeding individual, except some evidence that a slight reduction in the number of oaks within 75m predicted occupation by a local recruit (OR = 0.804, 95% CIs = 0.687–0.943, p = 0.007) or a specifically a female local recruit (OR = 0.769, 95% CIs = 0.617–0.959, p = 0.020). Greater EDI values (further from the edge) significantly predicted occupation by local recruits across all individuals (OR = 1.558, 95% CIs = 1.295–1.874, $p < 0.001$) and females (OR = 1.484, 95% CIs = 1.151–1.913, p = 0.002), but not males (OR = 1.098, 95% CIs = 0.851–1.416, p = 0.472). There was little evidence that any territory quality attributes predict the settled status of breeding individuals (Fig. S7; Table S2), except territories with a greater long-term popularity index were more likely to be occupied by settled males (OR = 1.075, 95% CIs = 1.024–1.128, p = 0.003), and territories with a higher EDI value were more likely to be acquired by settled individuals (OR = 1.047, 95% CIs = 1.005–1.089, p = 0.026).

##### 2.2 Spatial structure in residency status

There is weak evidence for spatial structure in immigration status across breeding territories when considering the whole population (median r assortativity coefficient = 0.007; 51% of annual standard errors overlap zero), females (median r = 0.029; 44% SEs overlap zero), or males (median r = -0.005; 47% SEs overlap zero). Similar results are found for settled status when considering the entire population (median r = 0.008; 51% SEs overlap zero), females (median r = 0.007; 38% SEs overlap zero) or males (median r = -0.019; 44% SEs overlap zero; Table S4 for all results).

###### *2.2.1 Temporal repeatability of spatial structure in residency status*

There was low repeatability in the residency status composition of spatial regions across years at all assessed spatial scales 25–200 ha (immigration status ICC range: 0.076–0.173; settled status ICC range: 0.024–0.062; Table S5).

##### 2.3 Spatial age structure as a continuous and discrete trait

As with binary age, we generally found weak positive age assortment across breeding territories when treating age as a discrete variable with five categories (1, 2, 3, 4, and 5 or more years-old) or as a continuous trait, whether considering the whole population (median discrete age r = 0.004, 58% SEs overlap zero; median continuous age r = 0.004, 58% SEs overlap zero), females (median discrete age r = -0.007, 60% SEs overlap zero; median continuous age r = -0.002, 53% SEs overlap zero), or males (median discrete age r = -0.013, 36% SEs overlap zero; median continuous age r = -0.012, 36% SEs overlap zero; Table S4 for all results).

### 3 Supplementary discussion

#### 3.1 Residency status and territory quality

We reveal some covariance between territory quality and residency status. Specifically, locally-hatched females are more likely to occupy territories at lower densities and those that are further from the edge, but this effect is not present in males. There is no association between the long-term popularity of a territory and the immigration status of the occupant in either sex, while there is weak evidence that immigrants may occupy territories with higher oak abundance. Interestingly, no territory quality attributes predict immigration status when considering males on their own. However, individuals that have newly dispersed into the woods are slightly more likely to occupy territories closer to the edge and newly immigrating males are very weakly associated with territories that have a lower long-term popularity index, while the settled status of females is not predicted by any territory quality attributes.

The strongest effects of lower territory density and greater distance from edge in predicting occupation by residents rather than immigrants, independent of age, might relate to greater site familiarity in locally-hatched individuals. Independent of the social advantages of high breeding density among adults as discussed in the main text, sites at lower density can provide better quality habitats if associated with greater foraging area (Wilkin et al. 2006), and edge sites often provide worse quality habitats due to overcrowding, increased predation and reduced foraging area (Paton 1994; Huhta et al. 1999; Batáry and Báldi 2004; Deng and Gao 2005; Wilkin et al. 2007a). Thus, residents with greater site familiarity (Krebs 1982; Wilkin et al. 2007a; Slagsvold and Wiebe 2018) and higher competitive ability (Sandell and Smith 1991) might be dominant over immigrants who settle in unoccupied areas of lower quality habitat. However, in this scenario we might expect relatively high assortment in how immigrants are spatially structured during breeding (Farine and Sheldon 2015, but see later discussion).

The effect of territory density and edge distance in predicting immigration status is not present when only assessing males, as with the other measures of territory quality. This may be related to sex-differences in dispersal dynamics in great tits, where females generally immigrate into sink populations such as Wytham from greater distances (Greenwood et al. 1979; Verhulst et al. 1997) and later in the year (Dhondt 1979; Hudde 1995; Verhulst et al. 1997). Therefore, immigrant males may be more familiar with the local environment compared to females, which is important given that males control initial territory acquisition (Krebs 1971, 1977; Harvey et al. 1979; Greenwood 1980; Clarke et al. 1997), thus potentially weakening the association between immigration status and territory quality in males compared to females. However, what then remains puzzling is the finding that new males (i.e. birds that have dispersed into the woods and are breeding for the first time) are more likely to occupy territories closer to the edge and those that have been occupied fewer times in the long-term. It would make sense that new males, and indeed all new individuals, are found closer to the edge considering that they might be expected to settle closer to their natal habitats, which are outside the main woodland (Wilkin et al. 2007a). However, by definition, all new birds are immigrants, but in their first year of breeding. Thus, our results suggest that the prediction that immigrants will occupy territories closer to the edge persists throughout lifespan when considering all individuals across the population, but this effect dissolves in males past the first breeding attempt. Related to discussion in the main text, this may be related to sex-differences in territory acquisition roles, where males initially acquire territories as the primary resource for obtaining a mate, whereas females assess and choose among males and their defended resources/territories (Krebs 1977; Greenwood 1980; Clarke et al. 1997). It might then be that immigrant males gain experience of the woods over time and move further from the edge past their first attempt to better quality territories that are less affected by negative edge effects (Wilkin et al. 2007a). This might also be mirrored in our results showing that new immigrant males are more likely to occupy territories which are less popular in the long-term, but this effect is not present when considering immigration status across lifespan.

#### 3.2 Spatial structure in residency status

We find similar results for spatial structure in immigration and settled status as with age structure, with a general pattern of slightly positive assortment, but still very weak. This might be indicative of the fact that despite some covariance in territory quality and these two traits, the association is not strong enough to generate overall spatial structure. Previous research in this population shows that immigration status is positively assorted across winter social behaviour both in terms of binary immigrant/resident status (Farine et al. 2015) and also arrival time into the woods (Farine and Sheldon 2015), but that this is mediated by spatial sorting rather than social decisions within flocks. Seeing as residency is an important determinant of dominance in great tits (Sandell and Smith 1991), this likely represents a scenario where resident birds exclude immigrants from high-quality areas of winter foraging, or immigrants purposefully associate in areas of lower quality to reduce chances of future exclusion from breeding territories that are already well populated by residents (Farine and Sheldon 2015; Farine et al. 2015). However, the results of our analyses reveal that there is not strong evidence that immigrants are strongly partitioned into areas of lower quality.

### 4 Supplementary figures

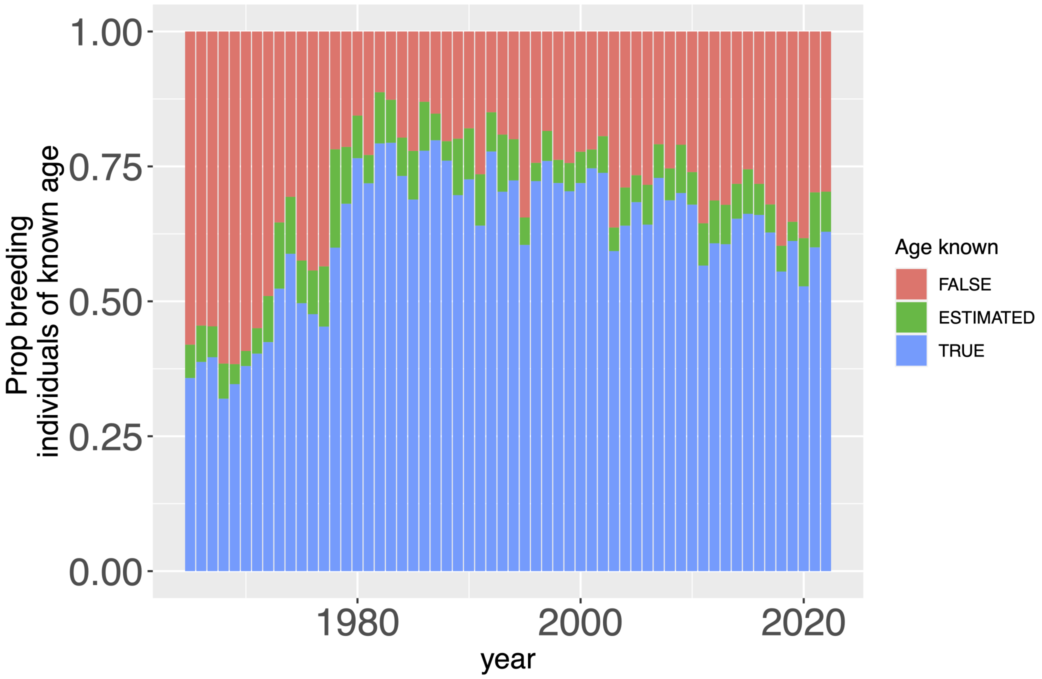

Fig. S1 - The annual proportion of breeding individuals where age is either unknown (individual not identified), estimated (individual first caught as an adult not in their first-year of life), or known (individual either first marked as a chick, or caught in their first-year of life) 1965–2022.

---

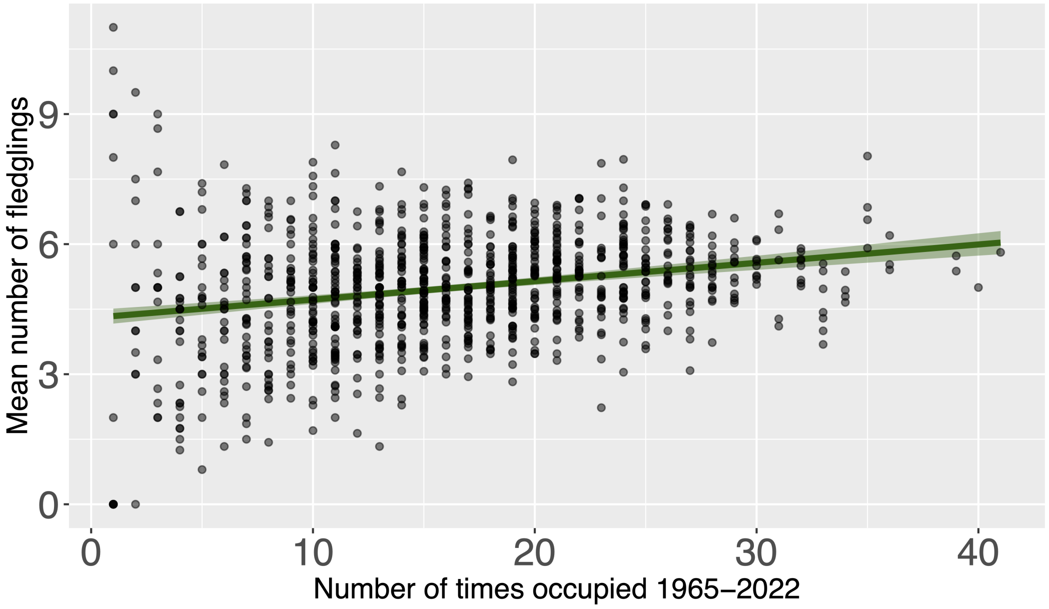

Fig. S2 - Relationship between the number of times a nest-box has been occupied 1965–2022 and the average number of chicks fledged from the box per breeding attempt. There is strong evidence for a weak to moderate relationship between these two variables (Spearman's correlation: r = 0.258, $p < 0.001$), therefore the frequency of occupation may provide a useful measure of quality.

---

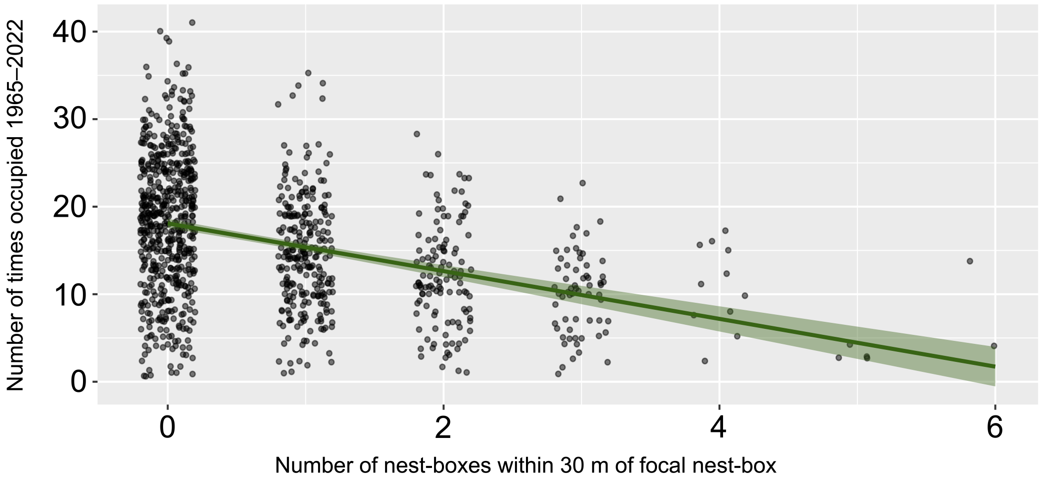

Fig. S3 - Relationship between the number of nest-boxes within a 30m radius of a focal box and the number of times it has been occupied 1965–2022. There is strong evidence for a weak to moderate negative relationship between these two variables (Spearman's correlation: r = -0.248, $p < 0.001$), therefore taking the residuals of a linear model between these two variables corrects for nest-box placing when calculating an index for long-term nest-box popularity. Points are jittered by 0.2 along the x-axis to avoid overplotting.

---

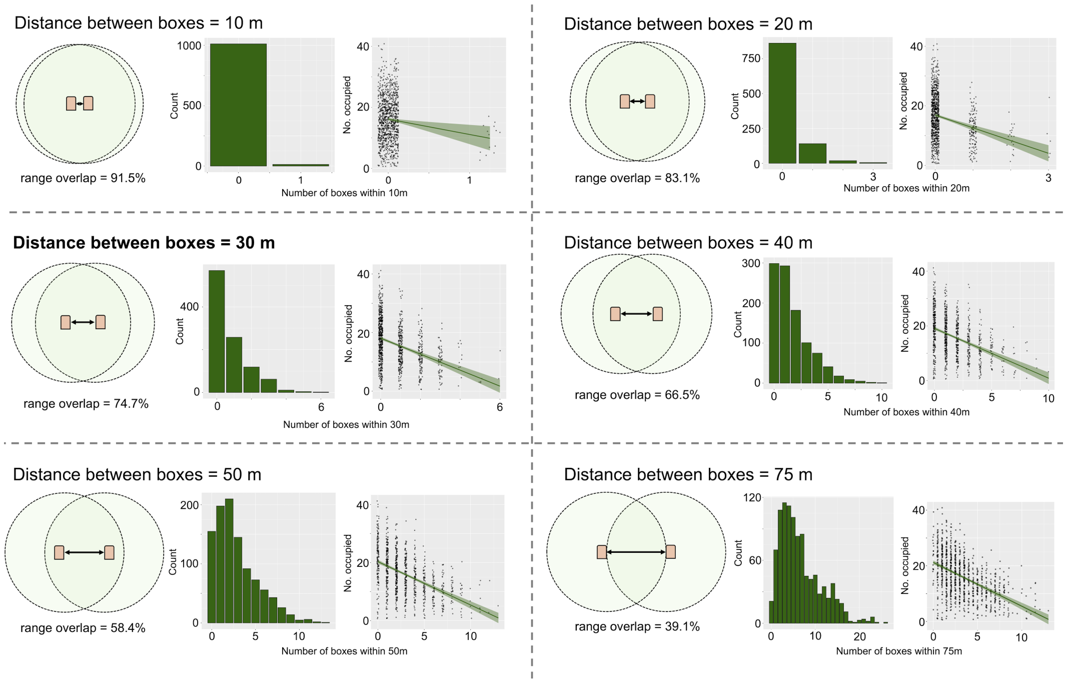

Fig. S4 - Overview of the different distances (10, 20, 30, 40, 50 and 75m) between a focal nest-box and neighbouring nest-boxes where, when placed at such densities, inter-annual switches between such boxes might result in largely similar territories regardless of which specific nest-box is occupied. For each focal distance: the Venn diagram displays the territorial range overlap (where territorial range is 75m); the histogram shows the number of nest-boxes across the study site which have a certain number of neighbouring boxes (on the x-axis) within the focal distance; and the plot on the right shows the relationship between the number of neighbouring boxes within the focal distance of a nest-box and the number of times said box has been occupied 1965–2022 (the green line indicates a linear regression between these two variables, the shading represents the 95% confidence intervals, and points are jittered by 0.1 along the x-axis to avoid overplotting).

---

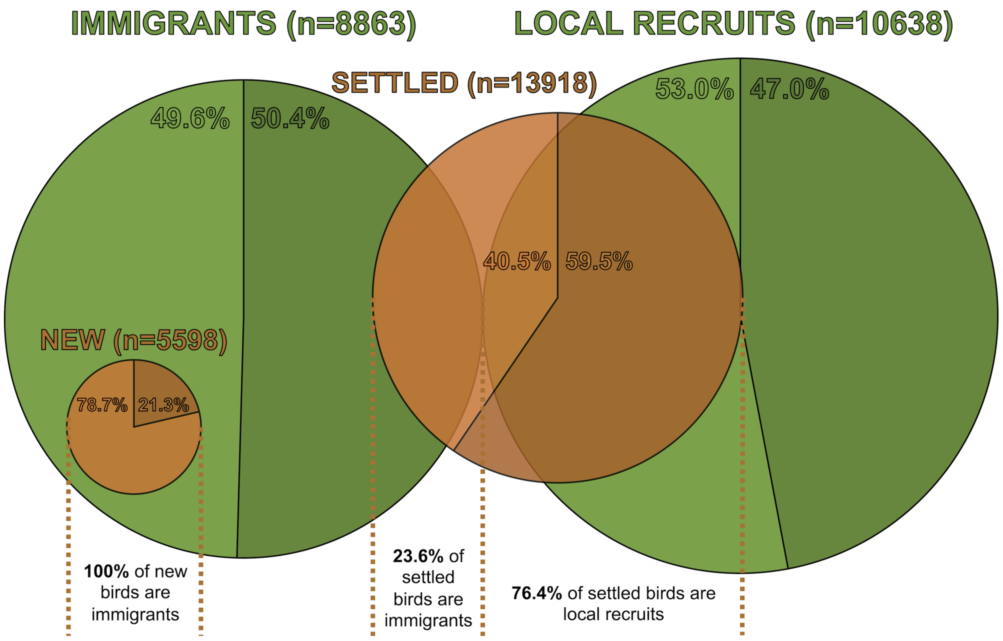

Fig. S5 - Synthesis of how different defined binary cohorts within the great tit population relate to one another. In all four pie charts, lighter colours represent the proportion of juveniles (yearlings), while the darker colours represent the proportion of adults (older than yearlings), with the proportion of each cohort fitting into these two age-classes expressed as a percentage in text. The green pie charts represent immigration status, where the two cohorts consist either of immigrants (hatched outside of Wytham Woods, n = 8863) or local recruits (hatched within Wytham, n = 10638). The two green pies' diameter is relative to the number of individuals found in these two cohorts. The orange pies represent settled status, where birds are either new (have arrived in Wytham within the last year, n = 5598) or settled (either first-year local recruits that are breeding for the first time, or individuals breeding for at least the second time that have previously bred in Wytham, n = 13918). The two orange pies' diameter is also relative to the number of individuals found in these two cohorts, but is not consistent with the scale used in the green pies. The overlap of orange with green pies represents the proportion of the respective settled status cohort that exists within the immigration status cohort (i.e. the entire orange pie representing new birds is within the green immigrants' pie as all new birds are immigrants, whereas there is only minimal overlap between the orange pie representing settled birds and the green immigrants pie as only 23.6% of settled birds originally immigrated into Wytham).

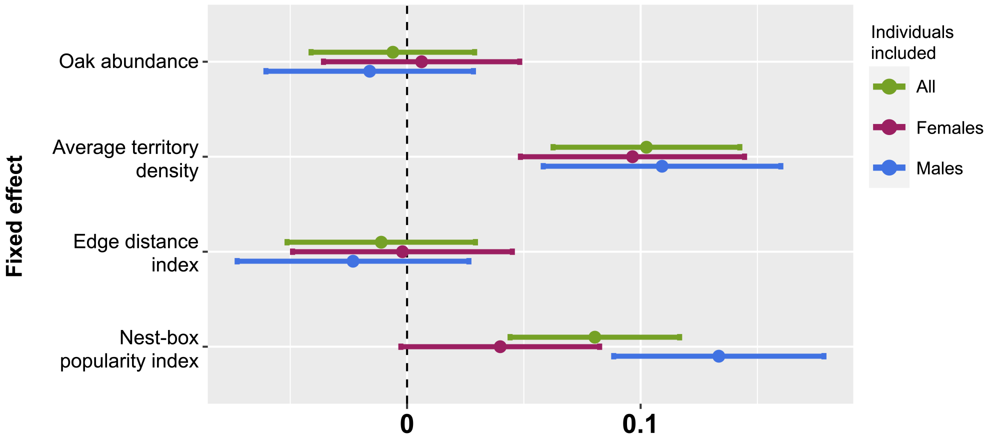

Fig. S6 - Point estimates of odds ratios obtained from generalised linear mixed-effects models. Each point corresponds to a specific level of the fixed effects (on the y-axis), and error bars denote 95% confidence intervals. The analysis examines the association between the four measures of territory quality on the odds of the breeding individual being a local recruit. Green points are from analysis assessing all individuals, purple are only females, and blue are only males.

---

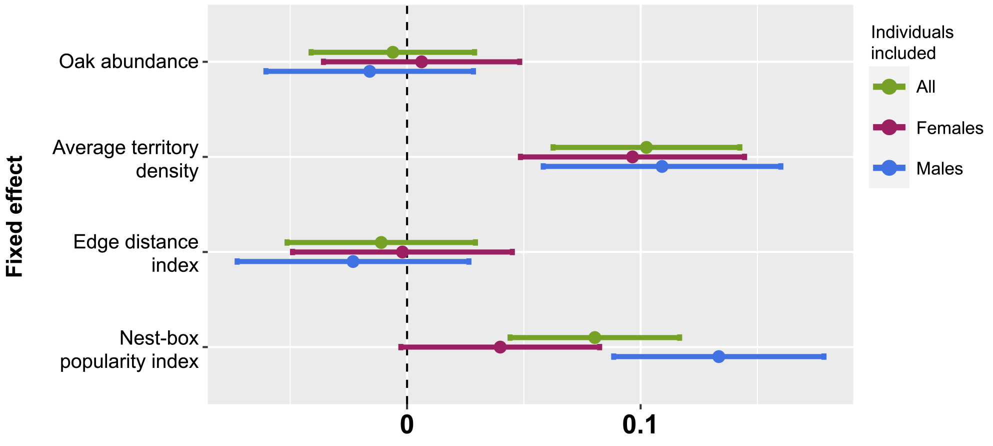

Fig. S7 - Point estimates of odds ratios obtained from generalised linear mixed-effects models. Each point corresponds to a specific level of the fixed effects (on the y-axis), and error bars denote 95% confidence intervals. The analysis examines the association between the four measures of territory quality on the odds of the breeding individual being a settled bird. Green points are from analysis assessing all individuals, purple are only females, and blue are only males.

---

#### 5 Supplementary tables

Table S1 - Binomial generalised linear mixed-effects model structure. Each model was run with all individuals; only females; and only males.

| Response variables | Fixed effects | Random effects |
|---|---|---|
| Age (0/1 = juvenile/adult) | Oak abundance | Individual ID |
| Immigration status (0/1 = immigrant/local recruit) | Average territory density | Nest-box ID |
| Settled status (0/1 = new/settled) | Edge distance index | Year |
| | Nest-box popularity index | |

Table S2 - Binomial generalised linear mixed-effects model results for predictors of age, immigration status and settled status, for all individuals, females and males separately, with individual ID, nest-box ID, and year included as random effects. Cells with light green shading indicate significant test results where p value $< 0.05$.

| Response variable | Individuals included | Fixed effect | $\beta$ | Odds ratio | 95% confidence intervals | p value |
|---|---|---|---|---|---|---|
| Age | All | Oak abundance | -0.006 | 0.994 | 0.960–1.029 | 0.734 |
| | | Average territory density | 0.102 | 1.108 | 1.065–1.153 | <0.001 |
| | | Edge distance index | -0.011 | 0.989 | 0.950–1.030 | 0.591 |
| | | Nest-box popularity index | 0.080 | 1.084 | 1.045–1.124 | <0.001 |
| | Females | Oak abundance | 0.006 | 1.006 | 0.965–1.049 | 0.771 |
| | | Average territory density | 0.097 | 1.101 | 1.050–1.155 | <0.001 |
| | | Edge distance index | -0.002 | 0.998 | 0.952–1.046 | 0.935 |
| | | Nest-box popularity index | 0.040 | 1.041 | 0.997–1.086 | 0.066 |
| | Males | Oak abundance | -0.016 | 0.984 | 0.941–1.029 | 0.480 |
| | | Average territory density | 0.109 | 1.115 | 1.060–1.174 | <0.001 |
| | | Edge distance index | -0.023 | 0.977 | 0.930–1.027 | 0.361 |
| | | Nest-box popularity index | 0.133 | 1.143 | 1.092–1.195 | <0.001 |
| Immigration status | All | Oak abundance | -0.218 | 0.804 | 0.687–0.943 | 0.007 |
| | | Average territory density | -0.413 | 0.662 | 0.567–0.773 | <0.001 |
| | | Edge distance index | 0.443 | 1.558 | 1.295–1.874 | <0.001 |
| | | Nest-box popularity index | 0.014 | 1.014 | 0.899–1.144 | 0.817 |
| | Females | Oak abundance | -0.263 | 0.769 | 0.617–0.959 | 0.020 |
| | | Average territory density | -0.471 | 0.624 | 0.504–0.773 | <0.001 |
| | | Edge distance index | 0.395 | 1.484 | 1.151–1.913 | 0.002 |
| | | Nest-box popularity index | -0.064 | 0.938 | 0.794–1.108 | 0.450 |
| | Males | Oak abundance | -0.008 | 0.992 | 0.796–1.237 | 0.945 |
| | | Average territory density | -0.050 | 0.951 | 0.748–1.210 | 0.683 |
| | | Edge distance index | 0.093 | 1.098 | 0.851–1.416 | 0.472 |
| | | Nest-box popularity index | 0.045 | 1.046 | 0.846–1.293 | 0.680 |
| Settled status | All | Oak abundance | -0.024 | 0.976 | 0.943–1.011 | 0.179 |
| | | Average territory density | 0.008 | 1.008 | 0.968–1.048 | 0.711 |
| | | Edge distance index | 0.045 | 1.047 | 1.005–1.089 | 0.026 |
| | | Nest-box popularity index | 0.027 | 1.028 | 0.992–1.065 | 0.132 |
| | Females | Oak abundance | -0.031 | 0.969 | 0.927–1.013 | 0.164 |
| | | Average territory density | -0.007 | 0.993 | 0.944–1.044 | 0.789 |
| | | Edge distance index | 0.039 | 1.040 | 0.989–1.093 | 0.129 |
| | | Nest-box popularity index | -0.012 | 0.988 | 0.945–1.033 | 0.600 |
| | Males | Oak abundance | -0.016 | 0.985 | 0.939–1.033 | 0.521 |
| | | Average territory density | 0.022 | 1.022 | 0.968–1.080 | 0.425 |
| | | Edge distance index | 0.053 | 1.055 | 0.999–1.114 | 0.054 |
| | | Nest-box popularity index | 0.072 | 1.075 | 1.024–1.128 | 0.003 |

Table S3 - Binomial generalised linear mixed-effects model results for predictors of age for all individuals. Nest-box popularity index was calculated by taking the residuals of a linear model between the number of boxes within 30m of a focal nest-box and the number of times occupied between 1965–2022 in the main text. Here, nest-box popularity index was calculated by alternatively taking the residuals of a linear model between the number of boxes within 10, 20, 40, 50 and 75m of a focal nest-box and the number of times occupied. Cells with light green shading indicate significant test results where p value ¡ 0.05. Bold text face indicates the results for the 30m threshold used in the main text.

| Threshold distance between boxes | Fixed effect | $\beta$ | Odds ratio | 95% confidence intervals | p value |
|---|---|---|---|---|---|
| 10m | Oak abundance | -0.005 | 0.995 | 0.959–1.030 | 0.761 |
| | Average territory density | 0.107 | 1.113 | 1.072–1.154 | <0.001 |
| | Edge distance index | -0.011 | 0.989 | 0.949–1.030 | 0.594 |
| | Nest-box popularity index | 0.066 | 1.069 | 1.031–1.106 | <0.001 |
| 20m | Oak abundance | -0.006 | 0.994 | 0.959–1.029 | 0.738 |
| | Average territory density | 0.105 | 1.110 | 1.070–1.151 | <0.001 |
| | Edge distance index | -0.011 | 0.989 | 0.948–1.029 | 0.581 |
| | Nest-box popularity index | 0.071 | 1.073 | 1.036–1.110 | <0.001 |
| **30m** | **Oak abundance** | **-0.006** | **0.994** | **0.960–1.029** | **0.734** |
| | **Average territory density** | **0.102** | **1.108** | **1.065–1.153** | <**0.001** |
| | **Edge distance index** | **-0.011** | **0.989** | **0.950–1.030** | **0.591** |
| | **Nest-box popularity index** | **0.080** | **1.084** | **1.045–1.1124** | <**0.001** |
| 40m | Oak abundance | -0.007 | 0.993 | 0.958–1.028 | 0.685 |
| | Average territory density | 0.100 | 1.105 | 1.066–1.145 | <0.001 |
| | Edge distance index | -0.011 | 0.989 | 0.949–1.030 | 0.604 |
| | Nest-box popularity index | 0.083 | 1.086 | 1.050–1.112 | <0.001 |
| 50m | Oak abundance | -0.008 | 0.992 | 0.957–1.027 | 0.651 |
| | Average territory density | 0.097 | 1.102 | 1.062–1.141 | <0.001 |
| | Edge distance index | -0.010 | 0.990 | 0.950–1.030 | 0.635 |
| | Nest-box popularity index | 0.088 | 1.092 | 1.057–1.128 | <0.001 |
| 75m | Oak abundance | -0.009 | 0.991 | 0.956–1.027 | 0.632 |
| | Average territory density | 0.094 | 1.099 | 1.059–1.138 | <0.001 |
| | Edge distance index | -0.011 | 0.989 | 0.949–1.030 | 0.605 |
| | Nest-box popularity index | 0.077 | 1.080 | 1.044–1.116 | <0.001 |

Table S4 – Annual network *r* assortativity results and standard error intervals across breeding territories for: binary, discrete, and continuous age; immigration status, and settled status (assessed either across all individuals, just females, or just males); number of oaks within 75m of nest-box, mean territory density, edge distance index, and long-term occupancy rate of the nest-box; clutch size, number of chicks, number of fledglings, and binary success. Cells with light green shading indicate significant positive assortment where standard error does not overlap with 0, and light red shading indicates significant negative assortment.

| Year | Focal trait type | Focal trait | Individuals | *r* | SE intervals |
|---|---|---|---|---|---|
| 1978 | Age | Binary age | All | -0.0247 | -0.056–0.007 |
| | | | Females | 0.0159 | -0.043–0.075 |
| | | | Males | -0.0736 | -0.142—0.005 |
| | | Discrete age | All | -0.0230 | -0.045—0.001 |
| | | | Females | 0.0268 | -0.015–0.068 |
| | | | Males | -0.0478 | -0.094—0.001 |
| | | Continuous age | All | -0.0347 | -0.062—0.007 |
| | | | Females | 0.0441 | -0.008–0.096 |
| | | | Males | -0.1502 | -0.202—0.098 |
| | Residency status | Immigration status | All | -0.0230 | -0.054–0.008 |
| | | | Females | -0.1012 | -0.16—0.042 |
| | | | Males | 0.0622 | -0.006–0.13 |
| | | Settled status | All | -0.0289 | -0.059–0.001 |
| | | | Females | -0.0922 | -0.151—0.034 |
| | | | Males | -0.0189 | -0.09–0.052 |
| | Territory quality | Oak abundance | NA | 0.7783 | 0.757–0.799 |

| Year | Focal trait type | Focal trait | Individuals | *r* | SE intervals |
|---|---|---|---|---|---|
| | | Average territory density | NA | 0.6523 | 0.626–0.678 |
| | | Edge distance index | NA | 0.7709 | 0.756–0.786 |
| | | Nest-box popularity index | NA | 0.3226 | 0.287–0.359 |
| | Reproductive | Clutch size | NA | -0.0072 | -0.049–0.035 |
| | | Number of chicks | NA | 0.0308 | -0.019–0.08 |
| | | Number of fledglings | NA | 0.0206 | -0.027–0.068 |
| | | Binary success | NA | -0.0604 | -0.091—0.03 |
| 1979 | Age | Binary age | All | 0.0513 | 0.02–0.083 |
| | | | Females | -0.0940 | -0.152—0.036 |
| | | | Males | -0.0085 | -0.073–0.056 |
| | | Discrete age | All | 0.0343 | 0.013–0.055 |
| | | | Females | -0.0902 | -0.121—0.059 |
| | | | Males | -0.0234 | -0.069–0.022 |
| | | Continuous age | All | 0.0471 | 0.016–0.078 |
| | | | Females | -0.0219 | -0.073–0.029 |
| | | | Males | 0.0548 | -0.016–0.125 |
| | Residency status | Immigration status | All | -0.0005 | -0.033–0.032 |
| | | | Females | 0.0196 | -0.041–0.081 |
| | | | Males | 0.0536 | -0.011–0.118 |
| | | Settled status | All | -0.0549 | -0.087—0.023 |
| | | | Females | -0.0068 | -0.065–0.052 |
| | | | Males | 0.0686 | -0.001–0.138 |
| | Territory quality | Oak abundance | NA | 0.7871 | 0.76–0.814 |
| | | Average territory density | NA | 0.7081 | 0.681–0.736 |
| | | Edge distance index | NA | 0.7163 | 0.698–0.735 |
| | | Nest-box popularity index | NA | 0.2655 | 0.227–0.304 |
| | Reproductive | Clutch size | NA | 0.0938 | 0.052–0.135 |
| | | Number of chicks | NA | -0.0150 | -0.049–0.019 |
| | | Number of fledglings | NA | -0.0351 | -0.078–0.008 |
| | | Binary success | NA | -0.0573 | -0.094—0.02 |
| 1980 | Age | Binary age | All | 0.0429 | 0.006–0.08 |
| | | | Females | 0.0156 | -0.05–0.081 |
| | | | Males | 0.0706 | -0.01–0.152 |
| | | Discrete age | All | 0.0434 | 0.018–0.069 |
| | | | Females | -0.0237 | -0.067–0.02 |
| | | | Males | 0.0878 | 0.025–0.151 |
| | | Continuous age | All | -0.0027 | -0.035–0.029 |
| | | | Females | -0.0051 | -0.056–0.046 |
| | | | Males | -0.0076 | -0.081–0.066 |
| | Residency status | Immigration status | All | 0.0349 | -0.001–0.07 |
| | | | Females | 0.0153 | -0.049–0.08 |
| | | | Males | -0.0820 | -0.161—0.003 |
| | | Settled status | All | 0.0104 | -0.026–0.047 |
| | | | Females | 0.0762 | 0.011–0.141 |
| | | | Males | -0.1042 | -0.183—0.025 |
| | Territory quality | Oak abundance | NA | 0.6970 | 0.668–0.726 |
| | | Average territory density | NA | 0.4360 | 0.399–0.473 |
| | | Edge distance index | NA | 0.7193 | 0.698–0.74 |
| | | Nest-box popularity index | NA | 0.2980 | 0.255–0.341 |
| | Reproductive | Clutch size | NA | -0.0180 | -0.055–0.019 |
| | | Number of chicks | NA | 0.2434 | 0.193–0.294 |
| | | Number of fledglings | NA | 0.2094 | 0.159–0.26 |
| | | Binary success | NA | 0.2822 | 0.231–0.333 |
| 1981 | Age | Binary age | All | -0.0304 | -0.056—0.005 |
| | | | Females | -0.0479 | -0.096–0 |
| | | | Males | 0.0541 | -0.005–0.113 |
| | | Discrete age | All | -0.0160 | -0.034–0.002 |
| | | | Females | -0.0094 | -0.046–0.027 |
| | | | Males | 0.0847 | 0.036–0.133 |

| Year | Focal trait type | Focal trait | Individuals | *r* | SE intervals |
|---|---|---|---|---|---|
| | | Continuous age | All | -0.0279 | -0.055—0.001 |
| | | | Females | -0.0094 | -0.065–0.046 |
| | | | Males | 0.1165 | 0.039–0.194 |
| | Residency status | Immigration status | All | 0.0195 | -0.006–0.045 |
| | | | Females | 0.1335 | 0.086–0.18 |
| | | | Males | -0.0056 | -0.062–0.05 |
| | | Settled status | All | -0.0390 | -0.064—0.014 |
| | | | Females | 0.0479 | 0–0.096 |
| | | | Males | 0.0434 | -0.018–0.104 |
| | Territory quality | Oak abundance | NA | 0.7781 | 0.763–0.794 |
| | | Average territory density | NA | 0.6394 | 0.61–0.669 |
| | | Edge distance index | NA | 0.8526 | 0.844–0.861 |
| | | Nest-box popularity index | NA | 0.2931 | 0.265–0.321 |
| | Reproductive | Clutch size | NA | 0.0447 | 0.014–0.075 |
| | | Number of chicks | NA | 0.0534 | 0.022–0.085 |
| | | Number of fledglings | NA | 0.0569 | 0.026–0.088 |
| | | Binary success | NA | 0.0728 | 0.041–0.105 |
| 1982 | Age | Binary age | All | 0.0415 | 0.017–0.066 |
| | | | Females | 0.0903 | 0.046–0.135 |
| | | | Males | 0.0503 | 0.001–0.1 |
| | | Discrete age | All | 0.0274 | 0.012–0.043 |
| | | | Females | 0.0487 | 0.019–0.079 |
| | | | Males | 0.0489 | 0.014–0.084 |
| | | Continuous age | All | 0.0016 | -0.019–0.023 |
| | | | Females | 0.0290 | -0.008–0.066 |
| | | | Males | 0.0622 | 0.014–0.111 |
| | Residency status | Immigration status | All | -0.0301 | -0.054—0.006 |
| | | | Females | 0.1412 | 0.097–0.186 |
| | | | Males | -0.0201 | -0.071–0.03 |
| | | Settled status | All | 0.0071 | -0.017–0.031 |
| | | | Females | 0.1559 | 0.108–0.204 |
| | | | Males | -0.0460 | -0.092—0.001 |
| | Territory quality | Oak abundance | NA | 0.7354 | 0.71–0.761 |
| | | Average territory density | NA | 0.6014 | 0.57–0.633 |
| | | Edge distance index | NA | 0.7634 | 0.748–0.779 |
| | | Nest-box popularity index | NA | 0.2106 | 0.174–0.247 |
| | Reproductive | Clutch size | NA | 0.1464 | 0.112–0.181 |
| | | Number of chicks | NA | 0.0012 | -0.038–0.04 |
| | | Number of fledglings | NA | -0.0224 | -0.059–0.014 |
| | | Binary success | NA | -0.0580 | -0.092—0.024 |
| 1983 | Age | Binary age | All | 0.0911 | 0.073–0.109 |
| | | | Females | -0.0316 | -0.065–0.002 |
| | | | Males | -0.0333 | -0.07–0.004 |
| | | Discrete age | All | 0.0618 | 0.049–0.074 |
| | | | Females | -0.0070 | -0.031–0.017 |
| | | | Males | -0.0337 | -0.057—0.011 |
| | | Continuous age | All | 0.0759 | 0.057–0.094 |
| | | | Females | -0.0219 | -0.052–0.008 |
| | | | Males | -0.0228 | -0.054–0.008 |
| | Residency status | Immigration status | All | -0.0224 | -0.04—0.004 |
| | | | Females | -0.0069 | -0.04–0.027 |
| | | | Males | -0.0383 | -0.077–0 |
| | | Settled status | All | 0.0238 | 0.006–0.042 |
| | | | Females | 0.0075 | -0.026–0.041 |
| | | | Males | -0.0766 | -0.115—0.038 |
| | Territory quality | Oak abundance | NA | 0.7889 | 0.776–0.802 |
| | | Average territory density | NA | 0.5873 | 0.556–0.619 |
| | | Edge distance index | NA | 0.8664 | 0.86–0.873 |
| | | Nest-box popularity index | NA | 0.2899 | 0.266–0.314 |

| Year | Focal trait type | Focal trait | Individuals | *r* | SE intervals |
|---|---|---|---|---|---|
| | Reproductive | Clutch size | NA | 0.1671 | 0.14–0.194 |
| | | Number of chicks | NA | 0.0931 | 0.065–0.121 |
| | | Number of fledglings | NA | 0.1219 | 0.092–0.152 |
| | | Binary success | NA | 0.1020 | 0.068–0.136 |
| 1984 | Age | Binary age | All | 0.0265 | -0.005–0.058 |
| | | | Females | -0.0490 | -0.104–0.006 |
| | | | Males | -0.0030 | -0.076–0.07 |
| | | Discrete age | All | 0.0111 | -0.009–0.031 |
| | | | Females | 0.0006 | -0.04–0.041 |
| | | | Males | 0.0161 | -0.035–0.067 |
| | | Continuous age | All | 0.0156 | -0.015–0.046 |
| | | | Females | -0.0978 | -0.14—0.056 |
| | | | Males | 0.0756 | 0–0.151 |
| | Residency status | Immigration status | All | -0.0177 | -0.05–0.014 |
| | | | Females | 0.0325 | -0.023–0.088 |
| | | | Males | -0.1831 | -0.244—0.122 |
| | | Settled status | All | -0.0445 | -0.077—0.012 |
| | | | Females | -0.0736 | -0.123—0.024 |
| | | | Males | -0.1622 | -0.202—0.122 |
| | Territory quality | Oak abundance | NA | 0.7746 | 0.759–0.79 |
| | | Average territory density | NA | 0.6586 | 0.637–0.68 |
| | | Edge distance index | NA | 0.7714 | 0.758–0.785 |
| | | Nest-box popularity index | NA | 0.1574 | 0.121–0.193 |
| | Reproductive | Clutch size | NA | 0.1128 | 0.078–0.147 |
| | | Number of chicks | NA | 0.1639 | 0.13–0.198 |
| | | Number of fledglings | NA | 0.1327 | 0.097–0.168 |
| | | Binary success | NA | 0.1139 | 0.077–0.151 |
| 1985 | Age | Binary age | All | 0.0081 | -0.021–0.037 |
| | | | Females | 0.0187 | -0.038–0.075 |
| | | | Males | -0.0562 | -0.114–0.002 |
| | | Discrete age | All | -0.0063 | -0.026–0.013 |
| | | | Females | 0.0057 | -0.036–0.048 |
| | | | Males | -0.0818 | -0.114—0.049 |
| | | Continuous age | All | -0.0191 | -0.046–0.008 |
| | | | Females | 0.0049 | -0.048–0.058 |
| | | | Males | -0.0406 | -0.088–0.007 |
| | Residency status | Immigration status | All | 0.0103 | -0.02–0.04 |
| | | | Females | 0.0745 | 0.016–0.133 |
| | | | Males | -0.1155 | -0.172—0.059 |
| | | Settled status | All | 0.0292 | -0.001–0.06 |
| | | | Females | 0.1209 | 0.062–0.18 |
| | | | Males | -0.0715 | -0.126—0.017 |
| | Territory quality | Oak abundance | NA | 0.6730 | 0.648–0.698 |
| | | Average territory density | NA | 0.7026 | 0.681–0.724 |
| | | Edge distance index | NA | 0.7621 | 0.748–0.777 |
| | | Nest-box popularity index | NA | 0.2792 | 0.242–0.317 |
| | Reproductive | Clutch size | NA | 0.0420 | 0.003–0.081 |
| | | Number of chicks | NA | 0.0453 | 0.005–0.086 |
| | | Number of fledglings | NA | -0.0321 | -0.068–0.004 |
| | | Binary success | NA | -0.0506 | -0.085—0.017 |
| 1986 | Age | Binary age | All | 0.0590 | 0.034–0.083 |
| | | | Females | 0.0627 | 0.012–0.113 |
| | | | Males | 0.0949 | 0.045–0.144 |
| | | Discrete age | All | 0.0440 | 0.028–0.06 |
| | | | Females | 0.0403 | 0.007–0.074 |
| | | | Males | 0.0644 | 0.032–0.097 |
| | | Continuous age | All | 0.0143 | -0.01–0.038 |
| | | | Females | 0.0883 | 0.032–0.145 |
| | | | Males | -0.0227 | -0.065–0.02 |

| Year | Focal trait type | Focal trait | Individuals | *r* | SE intervals |
|---|---|---|---|---|---|
| | Residency status | Immigration status | All | 0.0669 | 0.043–0.091 |
| | | | Females | 0.0504 | 0.002–0.099 |
| | | | Males | 0.0364 | -0.013–0.086 |
| | | Settled status | All | 0.0144 | -0.009–0.038 |
| | | | Females | 0.1035 | 0.055–0.152 |
| | | | Males | -0.0920 | -0.139—0.045 |
| | Territory quality | Oak abundance | NA | 0.7713 | 0.75–0.793 |
| | | Average territory density | NA | 0.6548 | 0.625–0.685 |
| | | Edge distance index | NA | 0.8038 | 0.791–0.817 |
| | | Nest-box popularity index | NA | 0.3357 | 0.304–0.368 |
| | Reproductive | Clutch size | NA | 0.0836 | 0.054–0.113 |
| | | Number of chicks | NA | 0.0646 | 0.032–0.097 |
| | | Number of fledglings | NA | 0.0938 | 0.06–0.128 |
| | | Binary success | NA | 0.0297 | -0.008–0.068 |
| 1987 | Age | Binary age | All | 0.0247 | 0–0.049 |
| | | | Females | 0.0016 | -0.042–0.045 |
| | | | Males | -0.1140 | -0.171—0.057 |
| | | Discrete age | All | 0.0416 | 0.023–0.06 |
| | | | Females | 0.0049 | -0.026–0.036 |
| | | | Males | -0.0959 | -0.134—0.058 |
| | | Continuous age | All | 0.0798 | 0.038–0.121 |
| | | | Females | -0.0191 | -0.063–0.025 |
| | | | Males | -0.1377 | -0.176—0.099 |
| | Residency status | Immigration status | All | 0.0282 | 0.004–0.053 |
| | | | Females | 0.1845 | 0.141–0.228 |
| | | | Males | -0.0516 | -0.108–0.005 |
| | | Settled status | All | -0.0057 | -0.029–0.018 |
| | | | Females | 0.0547 | -0.005–0.114 |
| | | | Males | -0.1820 | -0.218—0.146 |
| | Territory quality | Oak abundance | NA | 0.7816 | 0.764–0.799 |
| | | Average territory density | NA | 0.7054 | 0.689–0.722 |
| | | Edge distance index | NA | 0.8135 | 0.802–0.825 |
| | | Nest-box popularity index | NA | 0.2417 | 0.211–0.272 |
| | Reproductive | Clutch size | NA | 0.0751 | 0.044–0.106 |
| | | Number of chicks | NA | 0.1020 | 0.055–0.149 |
| | | Number of fledglings | NA | 0.0169 | -0.019–0.053 |
| | | Binary success | NA | 0.0267 | -0.013–0.067 |
| 1988 | Age | Binary age | All | 0.0285 | 0.005–0.052 |
| | | | Females | 0.0388 | -0.004–0.082 |
| | | | Males | -0.1148 | -0.162—0.067 |
| | | Discrete age | All | 0.0268 | 0.011–0.042 |
| | | | Females | 0.0115 | -0.015–0.038 |
| | | | Males | -0.0927 | -0.12—0.066 |
| | | Continuous age | All | 0.0206 | -0.008–0.049 |
| | | | Females | -0.0366 | -0.075–0.001 |
| | | | Males | -0.0747 | -0.116—0.033 |
| | Residency status | Immigration status | All | 0.0266 | 0.003–0.05 |
| | | | Females | 0.0409 | -0.002–0.084 |
| | | | Males | 0.0632 | 0.01–0.116 |
| | | Settled status | All | -0.0271 | -0.051—0.004 |
| | | | Females | -0.0620 | -0.103—0.021 |
| | | | Males | -0.0178 | -0.068–0.033 |
| | Territory quality | Oak abundance | NA | 0.8245 | 0.811–0.838 |
| | | Average territory density | NA | 0.5981 | 0.567–0.629 |
| | | Edge distance index | NA | 0.8207 | 0.81–0.832 |
| | | Nest-box popularity index | NA | 0.3522 | 0.328–0.377 |
| | Reproductive | Clutch size | NA | 0.0823 | 0.048–0.117 |
| | | Number of chicks | NA | 0.0505 | 0.022–0.08 |
| | | Number of fledglings | NA | 0.1359 | 0.106–0.166 |

| Year | Focal trait type | Focal trait | Individuals | *r* | SE intervals |
|---|---|---|---|---|---|
| | | Binary success | NA | 0.0599 | 0.026–0.094 |
| 1989 | Age | Binary age | All | -0.0126 | -0.039–0.013 |
| | | | Females | 0.0459 | 0.002–0.089 |
| | | | Males | 0.1221 | 0.064–0.18 |
| | | Discrete age | All | -0.0068 | -0.023–0.01 |
| | | | Females | 0.0587 | 0.028–0.089 |
| | | | Males | 0.0456 | 0.011–0.08 |
| | | Continuous age | All | -0.0223 | -0.045–0 |
| | | | Females | -0.0163 | -0.046–0.013 |
| | | | Males | 0.1862 | 0.141–0.231 |
| | Residency status | Immigration status | All | 0.0150 | -0.011–0.041 |
| | | | Females | 0.1168 | 0.072–0.161 |
| | | | Males | -0.1679 | -0.224—0.111 |
| | | Settled status | All | 0.0076 | -0.019–0.034 |
| | | | Females | 0.0677 | 0.024–0.112 |
| | | | Males | -0.0808 | -0.134—0.027 |
| | Territory quality | Oak abundance | NA | 0.7541 | 0.729–0.78 |
| | | Average territory density | NA | 0.6372 | 0.618–0.657 |
| | | Edge distance index | NA | 0.8271 | 0.816–0.838 |
| | | Nest-box popularity index | NA | 0.2444 | 0.214–0.275 |
| | Reproductive | Clutch size | NA | 0.0916 | 0.06–0.123 |
| | | Number of chicks | NA | 0.1124 | 0.073–0.152 |
| | | Number of fledglings | NA | 0.0049 | -0.031–0.041 |
| | | Binary success | NA | 0.0370 | -0.001–0.075 |
| 1990 | Age | Binary age | All | 0.0018 | -0.026–0.029 |
| | | | Females | 0.0114 | -0.044–0.067 |
| | | | Males | 0.0256 | -0.029–0.08 |
| | | Discrete age | All | 0.0140 | -0.003–0.031 |
| | | | Females | 0.0465 | 0.01–0.083 |
| | | | Males | -0.0128 | -0.044–0.018 |
| | | Continuous age | All | 0.0020 | -0.027–0.031 |
| | | | Females | 0.0577 | 0.007–0.109 |
| | | | Males | 0.0272 | -0.035–0.09 |
| | Residency status | Immigration status | All | -0.0206 | -0.048–0.007 |
| | | | Females | -0.0686 | -0.125—0.012 |
| | | | Males | 0.1045 | 0.05–0.159 |
| | | Settled status | All | -0.0037 | -0.032–0.024 |
| | | | Females | -0.0957 | -0.152—0.039 |
| | | | Males | 0.1242 | 0.061–0.187 |
| | Territory quality | Oak abundance | NA | 0.8162 | 0.796–0.836 |
| | | Average territory density | NA | 0.6706 | 0.632–0.709 |
| | | Edge distance index | NA | 0.7964 | 0.783–0.81 |
| | | Nest-box popularity index | NA | 0.4233 | 0.392–0.454 |
| | Reproductive | Clutch size | NA | 0.0399 | 0.007–0.073 |
| | | Number of chicks | NA | 0.0502 | 0.013–0.087 |
| | | Number of fledglings | NA | 0.0478 | 0.009–0.087 |
| | | Binary success | NA | 0.0606 | 0.02–0.101 |
| 1991 | Age | Binary age | All | 0.0024 | -0.043–0.047 |
| | | | Females | 0.0080 | -0.074–0.09 |
| | | | Males | 0.0412 | -0.064–0.147 |
| | | Discrete age | All | 0.0091 | -0.017–0.035 |
| | | | Females | -0.0159 | -0.063–0.032 |
| | | | Males | 0.0022 | -0.055–0.059 |
| | | Continuous age | All | 0.0608 | 0.016–0.106 |
| | | | Females | 0.0423 | -0.044–0.128 |
| | | | Males | -0.0065 | -0.104–0.091 |
| | Residency status | Immigration status | All | -0.0403 | -0.086–0.005 |
| | | | Females | 0.2058 | 0.119–0.292 |
| | | | Males | 0.0929 | -0.004–0.189 |

| Year | Focal trait type | Focal trait | Individuals | *r* | SE intervals |
|---|---|---|---|---|---|
| | | Settled status | All | 0.0717 | 0.015–0.129 |
| | | | Females | 0.1599 | 0.057–0.263 |
| | | | Males | 0.2198 | 0.095–0.344 |
| | Territory quality | Oak abundance | NA | 0.6776 | 0.643–0.713 |
| | | Average territory density | NA | 0.5689 | 0.523–0.615 |
| | | Edge distance index | NA | 0.5780 | 0.54–0.616 |
| | | Nest-box popularity index | NA | 0.1270 | 0.071–0.183 |
| | Reproductive | Clutch size | NA | -0.0391 | -0.094–0.016 |
| | | Number of chicks | NA | 0.1191 | 0.034–0.205 |
| | | Number of fledglings | NA | 0.1026 | 0.041–0.164 |
| | | Binary success | NA | 0.2290 | 0.165–0.293 |
| 1992 | Age | Binary age | All | -0.0082 | -0.035–0.019 |
| | | | Females | -0.0394 | -0.091–0.012 |
| | | | Males | -0.0893 | -0.145—0.034 |
| | | Discrete age | All | 0.0033 | -0.015–0.021 |
| | | | Females | -0.0020 | -0.045–0.041 |
| | | | Males | -0.0520 | -0.085—0.019 |
| | | Continuous age | All | -0.0118 | -0.038–0.015 |
| | | | Females | 0.0779 | 0.017–0.138 |
| | | | Males | -0.0517 | -0.102—0.001 |
| | Residency status | Immigration status | All | -0.0302 | -0.057—0.003 |
| | | | Females | -0.0617 | -0.114—0.009 |
| | | | Males | -0.0048 | -0.059–0.05 |
| | | Settled status | All | -0.0405 | -0.067—0.014 |
| | | | Females | -0.0259 | -0.079–0.027 |
| | | | Males | -0.0329 | -0.086–0.02 |
| | Territory quality | Oak abundance | NA | 0.6877 | 0.663–0.712 |
| | | Average territory density | NA | 0.5633 | 0.538–0.589 |
| | | Edge distance index | NA | 0.7447 | 0.729–0.76 |
| | | Nest-box popularity index | NA | 0.2762 | 0.242–0.311 |
| | Reproductive | Clutch size | NA | -0.0220 | -0.058–0.014 |
| | | Number of chicks | NA | 0.0550 | 0.003–0.107 |
| | | Number of fledglings | NA | 0.0750 | 0.03–0.12 |
| | | Binary success | NA | 0.0988 | 0.053–0.144 |
| 1993 | Age | Binary age | All | 0.0437 | 0.022–0.065 |
| | | | Females | 0.0260 | -0.014–0.066 |
| | | | Males | -0.0439 | -0.088–0 |
| | | Discrete age | All | 0.0364 | 0.02–0.053 |
| | | | Females | -0.0019 | -0.03–0.026 |
| | | | Males | -0.0452 | -0.074—0.016 |
| | | Continuous age | All | 0.0161 | -0.002–0.035 |
| | | | Females | 0.0145 | -0.023–0.052 |
| | | | Males | -0.0420 | -0.078—0.006 |
| | Residency status | Immigration status | All | 0.0580 | 0.037–0.079 |
| | | | Females | 0.0205 | -0.018–0.06 |
| | | | Males | -0.0111 | -0.056–0.034 |
| | | Settled status | All | 0.0551 | 0.034–0.076 |
| | | | Females | -0.0246 | -0.062–0.013 |
| | | | Males | 0.0490 | 0.004–0.094 |
| | Territory quality | Oak abundance | NA | 0.7931 | 0.774–0.812 |
| | | Average territory density | NA | 0.6147 | 0.592–0.637 |
| | | Edge distance index | NA | 0.8310 | 0.822–0.84 |
| | | Nest-box popularity index | NA | 0.3054 | 0.280–0.331 |
| | Reproductive | Clutch size | NA | 0.0716 | 0.044–0.1 |
| | | Number of chicks | NA | 0.0290 | -0.003–0.061 |
| | | Number of fledglings | NA | 0.0775 | 0.046–0.109 |
| | | Binary success | NA | 0.0297 | -0.001–0.061 |
| 1994 | Age | Binary age | All | 0.0158 | -0.005–0.037 |
| | | | Females | 0.0901 | 0.05–0.13 |

| Year | Focal trait type | Focal trait | Individuals | *r* | SE intervals |
|---|---|---|---|---|---|
| | | | Males | -0.0931 | -0.139—0.047 |
| | | Discrete age | All | 0.0189 | 0.004–0.034 |
| | | | Females | 0.0289 | -0.001–0.059 |
| | | | Males | -0.0614 | -0.091—0.032 |
| | | Continuous age | All | 0.0118 | -0.012–0.035 |
| | | | Females | 0.1016 | 0.068–0.136 |
| | | | Males | -0.0907 | -0.133—0.049 |
| | Residency status | Immigration status | All | 0.0026 | -0.019–0.024 |
| | | | Females | 0.0258 | -0.014–0.066 |
| | | | Males | 0.0639 | 0.018–0.11 |
| | | Settled status | All | 0.0074 | -0.014–0.029 |
| | | | Females | 0.0375 | -0.002–0.077 |
| | | | Males | 0.0276 | -0.015–0.071 |
| | Territory quality | Oak abundance | NA | 0.7981 | 0.784–0.812 |
| | | Average territory density | NA | 0.6940 | 0.677–0.711 |
| | | Edge distance index | NA | 0.8370 | 0.829–0.845 |
| | | Nest-box popularity index | NA | 0.2398 | 0.210–0.270 |
| | Reproductive | Clutch size | NA | 0.0600 | 0.026–0.094 |
| | | Number of chicks | NA | 0.0320 | -0.002–0.066 |
| | | Number of fledglings | NA | 0.0372 | 0.009–0.065 |
| | | Binary success | NA | -0.0026 | -0.031–0.026 |
| 1995 | Age | Binary age | All | 0.0201 | -0.01–0.05 |
| | | | Females | -0.0427 | -0.097–0.012 |
| | | | Males | -0.0521 | -0.113–0.009 |
| | | Discrete age | All | 0.0006 | -0.02–0.021 |
| | | | Females | -0.0178 | -0.055–0.02 |
| | | | Males | -0.0644 | -0.108—0.02 |
| | | Continuous age | All | -0.0196 | -0.047–0.008 |
| | | | Females | -0.0017 | -0.056–0.052 |
| | | | Males | 0.0037 | -0.053–0.06 |
| | Residency status | Immigration status | All | 0.0318 | 0.003–0.061 |
| | | | Females | -0.0564 | -0.111—0.002 |
| | | | Males | -0.0636 | -0.124—0.003 |
| | | Settled status | All | 0.0404 | 0.01–0.071 |
| | | | Females | -0.0255 | -0.08–0.029 |
| | | | Males | 0.0220 | -0.045–0.089 |
| | Territory quality | Oak abundance | NA | 0.7852 | 0.773–0.798 |
| | | Average territory density | NA | 0.6551 | 0.638–0.673 |
| | | Edge distance index | NA | 0.8579 | 0.852–0.864 |
| | | Nest-box popularity index | NA | 0.2155 | 0.191–0.240 |
| | Reproductive | Clutch size | NA | 0.0504 | 0.025–0.076 |
| | | Number of chicks | NA | 0.0043 | -0.022–0.03 |
| | | Number of fledglings | NA | 0.0564 | 0.031–0.082 |
| | | Binary success | NA | 0.0104 | -0.015–0.036 |
| 1996 | Age | Binary age | All | -0.0252 | -0.046—0.004 |
| | | | Females | -0.0880 | -0.125—0.051 |
| | | | Males | -0.1207 | -0.166—0.075 |
| | | Discrete age | All | -0.0114 | -0.025–0.002 |
| | | | Females | -0.0624 | -0.084—0.04 |
| | | | Males | -0.0471 | -0.078—0.016 |
| | | Continuous age | All | -0.0258 | -0.046—0.006 |
| | | | Females | -0.0339 | -0.07–0.002 |
| | | | Males | -0.0906 | -0.134—0.048 |
| | Residency status | Immigration status | All | -0.0066 | -0.027–0.014 |
| | | | Females | 0.0446 | 0.006–0.083 |
| | | | Males | 0.1154 | 0.065–0.165 |
| | | Settled status | All | 0.0086 | -0.012–0.029 |
| | | | Females | 0.0713 | 0.032–0.111 |
| | | | Males | 0.0296 | -0.018–0.077 |

| Year | Focal trait type | Focal trait | Individuals | *r* | SE intervals |
|---|---|---|---|---|---|
| | Territory quality | Oak abundance | NA | 0.7980 | 0.787–0.809 |
| | | Average territory density | NA | 0.5983 | 0.573–0.624 |
| | | Edge distance index | NA | 0.8811 | 0.876–0.886 |
| | | Nest-box popularity index | NA | 0.2756 | 0.253–0.298 |
| | Reproductive | Clutch size | NA | 0.2008 | 0.179–0.223 |
| | | Number of chicks | NA | 0.0867 | 0.062–0.112 |
| | | Number of fledglings | NA | 0.0155 | -0.009–0.04 |
| | | Binary success | NA | -0.0215 | -0.047–0.004 |
| 1997 | Age | Binary age | All | 0.0260 | 0.004–0.048 |
| | | | Females | -0.0296 | -0.07–0.011 |
| | | | Males | 0.0635 | 0.017–0.11 |
| | | Discrete age | All | 0.0130 | -0.002–0.028 |
| | | | Females | -0.0004 | -0.027–0.026 |
| | | | Males | 0.0432 | 0.017–0.069 |
| | | Continuous age | All | 0.0382 | 0.017–0.059 |
| | | | Females | -0.0792 | -0.113—0.046 |
| | | | Males | -0.0529 | -0.097—0.009 |
| | Residency status | Immigration status | All | 0.0387 | 0.017–0.061 |
| | | | Females | 0.0579 | 0.015–0.101 |
| | | | Males | 0.0434 | -0.004–0.091 |
| | | Settled status | All | 0.0365 | 0.013–0.06 |
| | | | Females | 0.1001 | 0.051–0.149 |
| | | | Males | -0.0660 | -0.103—0.029 |
| | Territory quality | Oak abundance | NA | 0.8111 | 0.799–0.823 |
| | | Average territory density | NA | 0.6738 | 0.653–0.694 |
| | | Edge distance index | NA | 0.8426 | 0.835–0.85 |
| | | Nest-box popularity index | NA | 0.3552 | 0.332–0.378 |
| | Reproductive | Clutch size | NA | 0.0502 | 0.019–0.081 |
| | | Number of chicks | NA | -0.0105 | -0.036–0.015 |
| | | Number of fledglings | NA | -0.0450 | -0.072—0.018 |
| | | Binary success | NA | -0.0309 | -0.055—0.007 |
| 1998 | Age | Binary age | All | -0.0499 | -0.073—0.027 |
| | | | Females | -0.0989 | -0.138—0.059 |
| | | | Males | 0.0632 | 0.009–0.117 |
| | | Discrete age | All | -0.0202 | -0.034—0.007 |
| | | | Females | -0.0728 | -0.096—0.05 |
| | | | Males | -0.0139 | -0.041–0.014 |
| | | Continuous age | All | -0.0836 | -0.104—0.063 |
| | | | Females | -0.0532 | -0.096—0.011 |
| | | | Males | 0.0635 | -0.012–0.139 |
| | Residency status | Immigration status | All | 0.0019 | -0.021–0.025 |
| | | | Females | -0.0105 | -0.051–0.031 |
| | | | Males | 0.0565 | 0.002–0.111 |
| | | Settled status | All | 0.0284 | 0.004–0.052 |
| | | | Females | -0.0266 | -0.064–0.011 |
| | | | Males | -0.0372 | -0.085–0.011 |
| | Territory quality | Oak abundance | NA | 0.7732 | 0.759–0.788 |
| | | Average territory density | NA | 0.6634 | 0.643–0.684 |
| | | Edge distance index | NA | 0.8631 | 0.856–0.87 |
| | | Nest-box popularity index | NA | 0.2128 | 0.188–0.238 |
| | Reproductive | Clutch size | NA | 0.0402 | 0.015–0.066 |
| | | Number of chicks | NA | -0.0023 | -0.028–0.024 |
| | | Number of fledglings | NA | 0.0387 | 0.012–0.066 |
| | | Binary success | NA | -0.0288 | -0.056—0.002 |
| 1999 | Age | Binary age | All | 0.0381 | 0.01–0.066 |
| | | | Females | 0.0090 | -0.041–0.059 |
| | | | Males | -0.0576 | -0.122–0.007 |
| | | Discrete age | All | 0.0170 | -0.001–0.035 |
| | | | Females | 0.0077 | -0.023–0.038 |

| Year | Focal trait type | Focal trait | Individuals | *r* | SE intervals |
|---|---|---|---|---|---|
| | | | Males | -0.0054 | -0.051–0.04 |
| | | Continuous age | All | 0.0405 | 0.014–0.067 |
| | | | Females | -0.0492 | -0.098—0.001 |
| | | | Males | 0.0838 | 0.026–0.141 |
| | Residency status | Immigration status | All | -0.0128 | -0.04–0.015 |
| | | | Females | 0.0617 | 0.012–0.111 |
| | | | Males | 0.0649 | -0.001–0.131 |
| | | Settled status | All | 0.0147 | -0.013–0.042 |
| | | | Females | 0.0528 | 0.003–0.103 |
| | | | Males | 0.0065 | -0.06–0.073 |
| | Territory quality | Oak abundance | NA | 0.7623 | 0.747–0.778 |
| | | Average territory density | NA | 0.6251 | 0.594–0.656 |
| | | Edge distance index | NA | 0.8166 | 0.807–0.826 |
| | | Nest-box popularity index | NA | 0.1891 | 0.158–0.220 |
| | Reproductive | Clutch size | NA | 0.1081 | 0.078–0.139 |
| | | Number of chicks | NA | 0.0489 | 0.017–0.081 |
| | | Number of fledglings | NA | -0.0049 | -0.037–0.027 |
| | | Binary success | NA | -0.0716 | -0.106—0.037 |
| 2000 | Age | Binary age | All | 0.0026 | -0.024–0.029 |
| | | | Females | 0.0015 | -0.045–0.048 |
| | | | Males | -0.0879 | -0.151—0.025 |
| | | Discrete age | All | -0.0122 | -0.028–0.004 |
| | | | Females | 0.0278 | -0.003–0.059 |
| | | | Males | -0.0996 | -0.138—0.062 |
| | | Continuous age | All | -0.0123 | -0.037–0.012 |
| | | | Females | -0.0161 | -0.059–0.027 |
| | | | Males | -0.0353 | -0.097–0.026 |
| | Residency status | Immigration status | All | -0.0175 | -0.044–0.009 |
| | | | Females | -0.0830 | -0.129—0.037 |
| | | | Males | -0.0628 | -0.127–0.001 |
| | | Settled status | All | 0.0595 | 0.032–0.087 |
| | | | Females | 0.0000 | -0.048–0.048 |
| | | | Males | -0.0847 | -0.144—0.025 |
| | Territory quality | Oak abundance | NA | 0.7968 | 0.782–0.811 |
| | | Average territory density | NA | 0.6278 | 0.602–0.654 |
| | | Edge distance index | NA | 0.7887 | 0.777–0.8 |
| | | Nest-box popularity index | NA | 0.2575 | 0.226–0.289 |
| | Reproductive | Clutch size | NA | 0.0686 | 0.03–0.107 |
| | | Number of chicks | NA | 0.0822 | 0.05–0.114 |
| | | Number of fledglings | NA | 0.1269 | 0.091–0.163 |
| | | Binary success | NA | 0.1001 | 0.061–0.139 |
| 2001 | Age | Binary age | All | 0.0202 | -0.005–0.046 |
| | | | Females | 0.0782 | 0.035–0.121 |
| | | | Males | -0.0572 | -0.125–0.01 |
| | | Discrete age | All | 0.0151 | -0.001–0.031 |
| | | | Females | 0.0141 | -0.011–0.039 |
| | | | Males | 0.0345 | -0.015–0.084 |
| | | Continuous age | All | 0.0161 | -0.009–0.041 |
| | | | Females | 0.1567 | 0.11–0.204 |
| | | | Males | -0.1855 | -0.229—0.142 |
| | Residency status | Immigration status | All | 0.0168 | -0.009–0.043 |
| | | | Females | 0.1097 | 0.068–0.152 |
| | | | Males | 0.0529 | -0.019–0.125 |
| | | Settled status | All | 0.0014 | -0.025–0.028 |
| | | | Females | 0.0607 | 0.02–0.101 |
| | | | Males | 0.1072 | 0.023–0.192 |
| | Territory quality | Oak abundance | NA | 0.8620 | 0.851–0.873 |
| | | Average territory density | NA | 0.6157 | 0.591–0.64 |
| | | Edge distance index | NA | 0.8234 | 0.814–0.833 |

| Year | Focal trait type | Focal trait | Individuals | *r* | SE intervals |
|---|---|---|---|---|---|
| | | Nest-box popularity index | NA | 0.1715 | 0.144–0.199 |
| | Reproductive | Clutch size | NA | 0.0831 | 0.045–0.121 |
| | | Number of chicks | NA | 0.0144 | -0.018–0.047 |
| | | Number of fledglings | NA | 0.0400 | 0.01–0.07 |
| | | Binary success | NA | -0.0192 | -0.052–0.013 |
| 2002 | Age | Binary age | All | 0.0088 | -0.014–0.032 |
| | | | Females | -0.0513 | -0.09—0.013 |
| | | | Males | -0.0038 | -0.055–0.047 |
| | | Discrete age | All | -0.0164 | -0.032—0.001 |
| | | | Females | -0.0553 | -0.08—0.031 |
| | | | Males | -0.0195 | -0.055–0.016 |
| | | Continuous age | All | 0.0825 | 0.056–0.109 |
| | | | Females | 0.1227 | 0.074–0.171 |
| | | | Males | 0.0935 | 0.045–0.142 |
| | Residency status | Immigration status | All | -0.0176 | -0.04–0.005 |
| | | | Females | 0.0514 | 0.012–0.091 |
| | | | Males | -0.0993 | -0.151—0.048 |
| | | Settled status | All | 0.0122 | -0.013–0.037 |
| | | | Females | -0.0351 | -0.069—0.002 |
| | | | Males | 0.0364 | -0.029–0.102 |
| | Territory quality | Oak abundance | NA | 0.8361 | 0.824–0.849 |
| | | Average territory density | NA | 0.6655 | 0.639–0.692 |
| | | Edge distance index | NA | 0.8406 | 0.832–0.849 |
| | | Nest-box popularity index | NA | 0.3003 | 0.276–0.325 |
| | Reproductive | Clutch size | NA | 0.0567 | 0.029–0.084 |
| | | Number of chicks | NA | 0.0004 | -0.025–0.026 |
| | | Number of fledglings | NA | 0.0456 | 0.017–0.074 |
| | | Binary success | NA | 0.0550 | 0.024–0.086 |
| 2003 | Age | Binary age | All | -0.0351 | -0.069—0.001 |
| | | | Females | 0.1460 | 0.099–0.194 |
| | | | Males | -0.0014 | -0.092–0.089 |
| | | Discrete age | All | -0.0175 | -0.038–0.003 |
| | | | Females | 0.0832 | 0.05–0.116 |
| | | | Males | -0.0585 | -0.111—0.006 |
| | | Continuous age | All | -0.0975 | -0.124—0.071 |
| | | | Females | 0.0787 | 0.034–0.124 |
| | | | Males | -0.0218 | -0.091–0.048 |
| | Residency status | Immigration status | All | -0.0567 | -0.089—0.024 |
| | | | Females | 0.0470 | -0.004–0.098 |
| | | | Males | -0.0386 | -0.127–0.049 |
| | | Settled status | All | -0.0617 | -0.094—0.029 |
| | | | Females | 0.1625 | 0.108–0.217 |
| | | | Males | -0.0014 | -0.071–0.069 |
| | Territory quality | Oak abundance | NA | 0.8597 | 0.848–0.871 |
| | | Average territory density | NA | 0.6143 | 0.583–0.646 |
| | | Edge distance index | NA | 0.8303 | 0.822–0.838 |
| | | Nest-box popularity index | NA | 0.3236 | 0.303–0.344 |
| | Reproductive | Clutch size | NA | 0.0579 | 0.034–0.082 |
| | | Number of chicks | NA | -0.0374 | -0.063—0.011 |
| | | Number of fledglings | NA | 0.0568 | 0.031–0.082 |
| | | Binary success | NA | -0.0193 | -0.045–0.007 |
| 2004 | Age | Binary age | All | 0.0028 | -0.024–0.029 |
| | | | Females | -0.1146 | -0.162—0.067 |
| | | | Males | 0.0816 | 0.018–0.145 |
| | | Discrete age | All | -0.0261 | -0.043—0.009 |
| | | | Females | -0.0660 | -0.096—0.035 |
| | | | Males | 0.0322 | -0.008–0.073 |
| | | Continuous age | All | 0.0137 | -0.011–0.038 |
| | | | Females | -0.0453 | -0.099–0.009 |

| Year | Focal trait type | Focal trait | Individuals | *r* | SE intervals |
|---|---|---|---|---|---|
| | | | Males | 0.0577 | 0.01–0.105 |
| | Residency status | Immigration status | All | -0.0345 | -0.062—0.007 |
| | | | Females | -0.1067 | -0.155—0.058 |
| | | | Males | -0.1118 | -0.168—0.056 |
| | | Settled status | All | 0.0245 | -0.002–0.051 |
| | | | Females | -0.0077 | -0.059–0.043 |
| | | | Males | -0.0119 | -0.07–0.047 |
| | Territory quality | Oak abundance | NA | 0.7954 | 0.78–0.81 |
| | | Average territory density | NA | 0.6033 | 0.581–0.626 |
| | | Edge distance index | NA | 0.8286 | 0.82–0.838 |
| | | Nest-box popularity index | NA | 0.3120 | 0.290–0.334 |
| | Reproductive | Clutch size | NA | 0.1715 | 0.141–0.202 |
| | | Number of chicks | NA | 0.0370 | 0.009–0.065 |
| | | Number of fledglings | NA | 0.0868 | 0.059–0.115 |
| | | Binary success | NA | 0.0633 | 0.033–0.094 |
| 2005 | Age | Binary age | All | 0.0144 | -0.006–0.035 |
| | | | Females | 0.0889 | 0.052–0.126 |
| | | | Males | -0.0847 | -0.129—0.04 |
| | | Discrete age | All | 0.0064 | -0.008–0.02 |
| | | | Females | 0.0720 | 0.046–0.098 |
| | | | Males | -0.0575 | -0.084—0.031 |
| | | Continuous age | All | -0.0032 | -0.022–0.015 |
| | | | Females | 0.0316 | 0.001–0.062 |
| | | | Males | -0.0952 | -0.13—0.06 |
| | Residency status | Immigration status | All | 0.0127 | -0.007–0.033 |
| | | | Females | 0.0631 | 0.027–0.1 |
| | | | Males | -0.0668 | -0.114—0.019 |
| | | Settled status | All | -0.0146 | -0.034–0.005 |
| | | | Females | 0.0861 | 0.049–0.123 |
| | | | Males | -0.0778 | -0.124—0.031 |
| | Territory quality | Oak abundance | NA | 0.8316 | 0.822–0.841 |
| | | Average territory density | NA | 0.6622 | 0.646–0.678 |
| | | Edge distance index | NA | 0.8660 | 0.86–0.872 |
| | | Nest-box popularity index | NA | 0.2309 | 0.209–0.252 |
| | Reproductive | Clutch size | NA | 0.1261 | 0.102–0.15 |
| | | Number of chicks | NA | 0.0866 | 0.062–0.111 |
| | | Number of fledglings | NA | 0.1424 | 0.12–0.165 |
| | | Binary success | NA | 0.0944 | 0.071–0.118 |
| 2006 | Age | Binary age | All | -0.0047 | -0.028–0.019 |
| | | | Females | -0.0487 | -0.09—0.007 |
| | | | Males | 0.0687 | 0.015–0.122 |
| | | Discrete age | All | -0.0041 | -0.02–0.012 |
| | | | Females | 0.0344 | 0.002–0.066 |
| | | | Males | 0.0371 | 0.005–0.069 |
| | | Continuous age | All | -0.0066 | -0.03–0.017 |
| | | | Females | -0.0378 | -0.088–0.012 |
| | | | Males | -0.0092 | -0.063–0.045 |
| | Residency status | Immigration status | All | -0.0392 | -0.063—0.016 |
| | | | Females | 0.0099 | -0.032–0.052 |
| | | | Males | -0.1358 | -0.189—0.083 |
| | | Settled status | All | -0.0417 | -0.065—0.019 |
| | | | Females | -0.0835 | -0.125—0.042 |
| | | | Males | -0.1245 | -0.174—0.075 |
| | Territory quality | Oak abundance | NA | 0.8257 | 0.815–0.836 |
| | | Average territory density | NA | 0.6453 | 0.616–0.674 |
| | | Edge distance index | NA | 0.8591 | 0.853–0.865 |
| | | Nest-box popularity index | NA | 0.2990 | 0.275–0.323 |
| | Reproductive | Clutch size | NA | 0.0477 | 0.021–0.074 |
| | | Number of chicks | NA | 0.0183 | -0.005–0.042 |

| Year | Focal trait type | Focal trait | Individuals | *r* | SE intervals |
|---|---|---|---|---|---|
| | | Number of fledglings | NA | 0.1089 | 0.083–0.134 |
| | | Binary success | NA | 0.0865 | 0.059–0.114 |
| 2007 | Age | Binary age | All | -0.0226 | -0.04—0.005 |
| | | | Females | -0.0136 | -0.047–0.019 |
| | | | Males | -0.0607 | -0.102—0.02 |
| | | Discrete age | All | -0.0062 | -0.018–0.006 |
| | | | Females | -0.0112 | -0.035–0.013 |
| | | | Males | -0.0124 | -0.039–0.014 |
| | | Continuous age | All | -0.0166 | -0.034–0.001 |
| | | | Females | -0.0039 | -0.033–0.025 |
| | | | Males | -0.0993 | -0.136—0.063 |
| | Residency status | Immigration status | All | -0.0296 | -0.047—0.012 |
| | | | Females | -0.0118 | -0.044–0.02 |
| | | | Males | 0.0603 | 0.02–0.101 |
| | | Settled status | All | -0.0459 | -0.063—0.028 |
| | | | Females | -0.0723 | -0.104—0.041 |
| | | | Males | 0.0424 | 0–0.085 |
| | Territory quality | Oak abundance | NA | 0.8105 | 0.801–0.82 |
| | | Average territory density | NA | 0.6211 | 0.597–0.645 |
| | | Edge distance index | NA | 0.8904 | 0.886–0.895 |
| | | Nest-box popularity index | NA | 0.2454 | 0.224–0.266 |
| | Reproductive | Clutch size | NA | 0.0588 | 0.036–0.081 |
| | | Number of chicks | NA | 0.0589 | 0.036–0.082 |
| | | Number of fledglings | NA | 0.1363 | 0.113–0.16 |
| | | Binary success | NA | 0.1090 | 0.084–0.134 |
| 2008 | Age | Binary age | All | 0.0620 | 0.041–0.083 |
| | | | Females | 0.0860 | 0.049–0.123 |
| | | | Males | 0.1014 | 0.048–0.155 |
| | | Discrete age | All | 0.0478 | 0.034–0.062 |
| | | | Females | 0.0517 | 0.026–0.077 |
| | | | Males | 0.0588 | 0.025–0.093 |
| | | Continuous age | All | 0.0233 | 0.002–0.045 |
| | | | Females | 0.0680 | 0.029–0.108 |
| | | | Males | 0.1391 | 0.086–0.192 |
| | Residency status | Immigration status | All | 0.0070 | -0.014–0.028 |
| | | | Females | 0.0274 | -0.009–0.064 |
| | | | Males | -0.0880 | -0.142—0.034 |
| | | Settled status | All | -0.0036 | -0.024–0.017 |
| | | | Females | 0.0232 | -0.014–0.06 |
| | | | Males | -0.0699 | -0.122—0.018 |
| | Territory quality | Oak abundance | NA | 0.8341 | 0.825–0.843 |
| | | Average territory density | NA | 0.6573 | 0.626–0.689 |
| | | Edge distance index | NA | 0.8791 | 0.874–0.884 |
| | | Nest-box popularity index | NA | 0.2312 | 0.207–0.255 |
| | Reproductive | Clutch size | NA | 0.0543 | 0.013–0.095 |
| | | Number of chicks | NA | 0.0450 | 0.01–0.08 |
| | | Number of fledglings | NA | 0.1267 | 0.102–0.151 |
| | | Binary success | NA | 0.1121 | 0.086–0.138 |
| 2009 | Age | Binary age | All | 0.0257 | 0.002–0.05 |
| | | | Females | 0.0110 | -0.029–0.051 |
| | | | Males | 0.0041 | -0.053–0.061 |
| | | Discrete age | All | 0.0077 | -0.007–0.022 |
| | | | Females | -0.0270 | -0.05—0.004 |
| | | | Males | 0.0036 | -0.031–0.038 |
| | | Continuous age | All | 0.0468 | 0.021–0.073 |
| | | | Females | -0.0090 | -0.047–0.029 |
| | | | Males | -0.1442 | -0.196—0.092 |
| | Residency status | Immigration status | All | 0.0006 | -0.023–0.024 |
| | | | Females | 0.1482 | 0.109–0.187 |

| Year | Focal trait type | Focal trait | Individuals | $r$ | SE intervals |
|---|---|---|---|---|---|
| | | | Males | 0.0658 | 0.01–0.122 |
| | | Settled status | All | 0.0203 | -0.004–0.044 |
| | | | Females | 0.0710 | 0.03–0.112 |
| | | | Males | 0.0022 | -0.055–0.059 |
| | Territory quality | Oak abundance | NA | 0.7973 | 0.781–0.814 |
| | | Average territory density | NA | 0.7075 | 0.692–0.723 |
| | | Edge distance index | NA | 0.8175 | 0.808–0.827 |
| | | Nest-box popularity index | NA | 0.2576 | 0.232–0.283 |
| | Reproductive | Clutch size | NA | 0.0611 | 0.019–0.103 |
| | | Number of chicks | NA | 0.0373 | 0.006–0.069 |
| | | Number of fledglings | NA | 0.0674 | 0.038–0.097 |
| | | Binary success | NA | 0.0336 | 0.002–0.065 |
| 2010 | Age | Binary age | All | 0.0097 | -0.014–0.033 |
| | | | Females | 0.0561 | 0.015–0.098 |
| | | | Males | 0.0030 | -0.057–0.063 |
| | | Discrete age | All | 0.0016 | -0.013–0.016 |
| | | | Females | 0.0441 | 0.016–0.072 |
| | | | Males | -0.0699 | -0.099—0.041 |
| | | Continuous age | All | 0.0039 | -0.021–0.029 |
| | | | Females | -0.0004 | -0.04–0.039 |
| | | | Males | 0.0347 | -0.023–0.093 |
| | Residency status | Immigration status | All | 0.0378 | 0.014–0.061 |
| | | | Females | -0.1072 | -0.149—0.065 |
| | | | Males | 0.0315 | -0.027–0.09 |
| | | Settled status | All | -0.0129 | -0.036–0.011 |
| | | | Females | -0.0959 | -0.136—0.056 |
| | | | Males | -0.0453 | -0.105–0.014 |
| | Territory quality | Oak abundance | NA | 0.8335 | 0.821–0.846 |
| | | Average territory density | NA | 0.6586 | 0.641–0.676 |
| | | Edge distance index | NA | 0.8159 | 0.808–0.824 |
| | | Nest-box popularity index | NA | 0.2606 | 0.238–0.283 |
| | Reproductive | Clutch size | NA | 0.0208 | -0.004–0.046 |
| | | Number of chicks | NA | 0.0371 | 0.008–0.066 |
| | | Number of fledglings | NA | 0.1605 | 0.136–0.186 |
| | | Binary success | NA | 0.0753 | 0.047–0.103 |
| 2011 | Age | Binary age | All | 0.0139 | -0.022–0.05 |
| | | | Females | -0.0516 | -0.112–0.009 |
| | | | Males | -0.2251 | -0.307—0.144 |
| | | Discrete age | All | 0.0005 | -0.027–0.028 |
| | | | Females | -0.0454 | -0.088—0.003 |
| | | | Males | -0.1563 | -0.206—0.106 |
| | | Continuous age | All | -0.0166 | -0.049–0.016 |
| | | | Females | -0.0993 | -0.137—0.061 |
| | | | Males | -0.1716 | -0.252—0.091 |
| | Residency status | Immigration status | All | -0.0129 | -0.049–0.023 |
| | | | Females | -0.0081 | -0.071–0.055 |
| | | | Males | 0.0861 | 0–0.172 |
| | | Settled status | All | 0.0689 | 0.032–0.105 |
| | | | Females | 0.1009 | 0.032–0.17 |
| | | | Males | -0.1335 | -0.215—0.052 |
| | Territory quality | Oak abundance | NA | 0.7725 | 0.756–0.789 |
| | | Average territory density | NA | 0.6448 | 0.614–0.676 |
| | | Edge distance index | NA | 0.8191 | 0.809–0.829 |
| | | Nest-box popularity index | NA | 0.1989 | 0.171–0.227 |
| | Reproductive | Clutch size | NA | 0.0116 | -0.022–0.046 |
| | | Number of chicks | NA | -0.0794 | -0.107—0.052 |
| | | Number of fledglings | NA | 0.0634 | 0.033–0.094 |
| | | Binary success | NA | 0.0859 | 0.054–0.118 |
| 2012 | Age | Binary age | All | 0.0319 | 0.005–0.059 |

| Year | Focal trait type | Focal trait | Individuals | *r* | SE intervals |
|---|---|---|---|---|---|
| | | | Females | 0.0094 | -0.03–0.049 |
| | | | Males | 0.1383 | 0.061–0.215 |
| | | Discrete age | All | 0.0037 | -0.013–0.021 |
| | | | Females | -0.0185 | -0.043–0.006 |
| | | | Males | 0.1078 | 0.057–0.159 |
| | | Continuous age | All | 0.0485 | 0.023–0.074 |
| | | | Females | 0.0344 | -0.002–0.071 |
| | | | Males | 0.0821 | 0.012–0.153 |
| | Residency status | Immigration status | All | 0.0274 | 0–0.055 |
| | | | Females | 0.0060 | -0.032–0.044 |
| | | | Males | 0.2227 | 0.149–0.297 |
| | | Settled status | All | 0.0481 | 0.021–0.075 |
| | | | Females | -0.0332 | -0.075–0.008 |
| | | | Males | 0.0244 | -0.053–0.102 |
| | Territory quality | Oak abundance | NA | 0.8142 | 0.802–0.827 |
| | | Average territory density | NA | 0.6362 | 0.61–0.662 |
| | | Edge distance index | NA | 0.8513 | 0.844–0.858 |
| | | Nest-box popularity index | NA | 0.2746 | 0.250–0.299 |
| | Reproductive | Clutch size | NA | 0.0003 | -0.024–0.025 |
| | | Number of chicks | NA | -0.0561 | -0.081—0.031 |
| | | Number of fledglings | NA | 0.0377 | 0.012–0.063 |
| | | Binary success | NA | 0.0304 | 0.004–0.057 |
| 2013 | Age | Binary age | All | 0.0588 | 0.022–0.095 |
| | | | Females | 0.0964 | 0.026–0.167 |
| | | | Males | -0.1308 | -0.168—0.093 |
| | | Discrete age | All | 0.0154 | -0.005–0.036 |
| | | | Females | -0.0151 | -0.051–0.021 |
| | | | Males | -0.0393 | -0.072—0.007 |
| | | Continuous age | All | 0.0811 | 0.049–0.114 |
| | | | Females | 0.1041 | 0.031–0.178 |
| | | | Males | 0.1262 | 0.058–0.194 |
| | Residency status | Immigration status | All | 0.1080 | 0.071–0.145 |
| | | | Females | 0.0706 | 0.002–0.139 |
| | | | Males | 0.3344 | 0.258–0.411 |
| | | Settled status | All | 0.0683 | 0.031–0.105 |
| | | | Females | -0.0583 | -0.128–0.011 |
| | | | Males | 0.1747 | 0.095–0.254 |
| | Territory quality | Oak abundance | NA | 0.7544 | 0.737–0.772 |
| | | Average territory density | NA | 0.5966 | 0.573–0.621 |
| | | Edge distance index | NA | 0.7228 | 0.708–0.738 |
| | | Nest-box popularity index | NA | 0.2183 | 0.188–0.249 |
| | Reproductive | Clutch size | NA | 0.0093 | -0.022–0.04 |
| | | Number of chicks | NA | 0.0471 | 0.014–0.08 |
| | | Number of fledglings | NA | 0.0990 | 0.068–0.13 |
| | | Binary success | NA | 0.0615 | 0.028–0.095 |
| 2014 | Age | Binary age | All | -0.0663 | -0.095—0.038 |
| | | | Females | -0.0246 | -0.077–0.028 |
| | | | Males | -0.1372 | -0.198—0.076 |
| | | Discrete age | All | -0.0398 | -0.055—0.025 |
| | | | Females | -0.0273 | -0.055–0.001 |
| | | | Males | -0.0167 | -0.056–0.023 |
| | | Continuous age | All | -0.0176 | -0.045–0.009 |
| | | | Females | 0.0758 | 0.028–0.123 |
| | | | Males | -0.0010 | -0.068–0.066 |
| | Residency status | Immigration status | All | 0.0761 | 0.048–0.104 |
| | | | Females | 0.0423 | -0.01–0.095 |
| | | | Males | 0.0048 | -0.058–0.068 |
| | | Settled status | All | -0.0190 | -0.049–0.011 |
| | | | Females | 0.0772 | 0.025–0.129 |

| Year | Focal trait type | Focal trait | Individuals | *r* | SE intervals |
|---|---|---|---|---|---|
| | | | Males | -0.1633 | -0.212—0.115 |
| | Territory quality | Oak abundance | NA | 0.8164 | 0.802–0.83 |
| | | Average territory density | NA | 0.6610 | 0.646–0.676 |
| | | Edge distance index | NA | 0.7477 | 0.735–0.76 |
| | | Nest-box popularity index | NA | 0.2316 | 0.206–0.257 |
| | Reproductive | Clutch size | NA | 0.1034 | 0.073–0.134 |
| | | Number of chicks | NA | 0.0395 | 0.012–0.067 |
| | | Number of fledglings | NA | 0.0177 | -0.01–0.045 |
| | | Binary success | NA | -0.0130 | -0.043–0.017 |
| 2015 | Age | Binary age | All | 0.0089 | -0.016–0.034 |
| | | | Females | -0.0628 | -0.11—0.016 |
| | | | Males | 0.1243 | 0.07–0.179 |
| | | Discrete age | All | -0.0010 | -0.017–0.015 |
| | | | Females | -0.0292 | -0.061–0.003 |
| | | | Males | 0.0416 | 0.007–0.077 |
| | | Continuous age | All | 0.0188 | -0.003–0.041 |
| | | | Females | -0.0484 | -0.099–0.002 |
| | | | Males | 0.0700 | 0.028–0.112 |
| | Residency status | Immigration status | All | 0.0128 | -0.013–0.038 |
| | | | Females | -0.1064 | -0.153—0.059 |
| | | | Males | -0.1063 | -0.162—0.051 |
| | | Settled status | All | 0.0010 | -0.025–0.027 |
| | | | Females | -0.0739 | -0.12—0.028 |
| | | | Males | -0.0742 | -0.126—0.023 |
| | Territory quality | Oak abundance | NA | 0.7805 | 0.764–0.797 |
| | | Average territory density | NA | 0.6561 | 0.635–0.678 |
| | | Edge distance index | NA | 0.8040 | 0.794–0.814 |
| | | Nest-box popularity index | NA | 0.2880 | 0.264–0.312 |
| | Reproductive | Clutch size | NA | 0.1328 | 0.108–0.158 |
| | | Number of chicks | NA | 0.0142 | -0.018–0.047 |
| | | Number of fledglings | NA | 0.2255 | 0.198–0.253 |
| | | Binary success | NA | 0.0534 | 0.022–0.085 |
| 2016 | Age | Binary age | All | -0.0484 | -0.078—0.019 |
| | | | Females | 0.0101 | -0.045–0.065 |
| | | | Males | 0.1141 | 0.046–0.182 |
| | | Discrete age | All | -0.0153 | -0.034–0.003 |
| | | | Females | -0.0246 | -0.057–0.008 |
| | | | Males | 0.0364 | -0.005–0.078 |
| | | Continuous age | All | -0.0135 | -0.043–0.016 |
| | | | Females | 0.0541 | 0.003–0.105 |
| | | | Males | -0.0696 | -0.117—0.022 |
| | Residency status | Immigration status | All | -0.0543 | -0.083—0.025 |
| | | | Females | -0.1133 | -0.168—0.059 |
| | | | Males | 0.0277 | -0.041–0.097 |
| | | Settled status | All | -0.0477 | -0.078—0.017 |
| | | | Females | -0.0203 | -0.073–0.033 |
| | | | Males | 0.0211 | -0.049–0.091 |
| | Territory quality | Oak abundance | NA | 0.7478 | 0.726–0.769 |
| | | Average territory density | NA | 0.6688 | 0.649–0.688 |
| | | Edge distance index | NA | 0.8021 | 0.79–0.814 |
| | | Nest-box popularity index | NA | 0.2097 | 0.178–0.241 |
| | Reproductive | Clutch size | NA | 0.1184 | 0.068–0.169 |
| | | Number of chicks | NA | 0.0529 | 0.016–0.09 |
| | | Number of fledglings | NA | 0.1876 | 0.157–0.218 |
| | | Binary success | NA | 0.1227 | 0.089–0.156 |
| 2017 | Age | Binary age | All | 0.0289 | -0.004–0.061 |
| | | | Females | -0.0404 | -0.1–0.02 |
| | | | Males | -0.0637 | -0.127—0.001 |
| | | Discrete age | All | 0.0181 | 0–0.036 |

| Year | Focal trait type | Focal trait | Individuals | $r$ | SE intervals |
|---|---|---|---|---|---|
| | | | Females | -0.0186 | -0.054–0.016 |
| | | | Males | -0.0124 | -0.052–0.027 |
| | | Continuous age | All | 0.0395 | 0.006–0.072 |
| | | | Females | -0.0209 | -0.075–0.033 |
| | | | Males | -0.0124 | -0.083–0.058 |
| | Residency status | Immigration status | All | 0.0607 | 0.029–0.093 |
| | | | Females | 0.0602 | -0.001–0.121 |
| | | | Males | 0.0856 | 0.018–0.153 |
| | | Settled status | All | 0.0359 | 0.002–0.07 |
| | | | Females | -0.0499 | -0.109–0.009 |
| | | | Males | -0.0733 | -0.128—0.019 |
| | Territory quality | Oak abundance | NA | 0.7128 | 0.695–0.73 |
| | | Average territory density | NA | 0.6461 | 0.624–0.668 |
| | | Edge distance index | NA | 0.7817 | 0.77–0.793 |
| | | Nest-box popularity index | NA | 0.2846 | 0.257–0.312 |
| | Reproductive | Clutch size | NA | 0.0744 | 0.042–0.107 |
| | | Number of chicks | NA | 0.1504 | 0.113–0.187 |
| | | Number of fledglings | NA | 0.1988 | 0.168–0.229 |
| | | Binary success | NA | 0.1856 | 0.152–0.219 |
| 2018 | Age | Binary age | All | 0.0245 | -0.018–0.067 |
| | | | Females | -0.0260 | -0.1–0.048 |
| | | | Males | 0.0418 | -0.085–0.168 |
| | | Discrete age | All | 0.0475 | 0.019–0.076 |
| | | | Females | -0.0779 | -0.118—0.038 |
| | | | Males | 0.0508 | -0.022–0.123 |
| | | Continuous age | All | -0.0142 | -0.06–0.032 |
| | | | Females | 0.2417 | 0.141–0.342 |
| | | | Males | -0.0483 | -0.179–0.083 |
| | Residency status | Immigration status | All | 0.0373 | -0.005–0.079 |
| | | | Females | 0.0290 | -0.048–0.106 |
| | | | Males | -0.0951 | -0.212–0.021 |
| | | Settled status | All | 0.0195 | -0.022–0.061 |
| | | | Females | -0.0155 | -0.089–0.058 |
| | | | Males | 0.0117 | -0.116–0.14 |
| | Territory quality | Oak abundance | NA | 0.7464 | 0.725–0.767 |
| | | Average territory density | NA | 0.6708 | 0.651–0.691 |
| | | Edge distance index | NA | 0.7854 | 0.774–0.797 |
| | | Nest-box popularity index | NA | 0.2077 | 0.180–0.236 |
| | Reproductive | Clutch size | NA | 0.0436 | 0.016–0.071 |
| | | Number of chicks | NA | 0.1792 | 0.143–0.216 |
| | | Number of fledglings | NA | 0.1574 | 0.123–0.192 |
| | | Binary success | NA | 0.1641 | 0.124–0.204 |
| 2019 | Age | Binary age | All | 0.0106 | -0.02–0.041 |
| | | | Females | -0.0334 | -0.095–0.028 |
| | | | Males | 0.0261 | -0.035–0.087 |
| | | Discrete age | All | 0.0028 | -0.018–0.024 |
| | | | Females | -0.0196 | -0.058–0.019 |
| | | | Males | 0.0154 | -0.026–0.057 |
| | | Continuous age | All | 0.0404 | 0.012–0.069 |
| | | | Females | -0.0912 | -0.137—0.046 |
| | | | Males | 0.1849 | 0.098–0.272 |
| | Residency status | Immigration status | All | 0.0515 | 0.021–0.082 |
| | | | Females | 0.1250 | 0.065–0.185 |
| | | | Males | -0.0398 | -0.104–0.024 |
| | | Settled status | All | 0.0497 | 0.019–0.08 |
| | | | Females | 0.0882 | 0.027–0.149 |
| | | | Males | 0.0065 | -0.058–0.071 |
| | Territory quality | Oak abundance | NA | 0.7684 | 0.756–0.781 |
| | | Average territory density | NA | 0.6372 | 0.618–0.656 |

| Year | Focal trait type | Focal trait | Individuals | *r* | SE intervals |
|---|---|---|---|---|---|
| | | Edge distance index | NA | 0.8197 | 0.811–0.829 |
| | | Nest-box popularity index | NA | 0.2097 | 0.184–0.236 |
| | Reproductive | Clutch size | NA | 0.0154 | -0.01–0.041 |
| | | Number of chicks | NA | 0.0465 | 0.019–0.074 |
| | | Number of fledglings | NA | 0.0219 | -0.003–0.047 |
| | | Binary success | NA | 0.0068 | -0.018–0.032 |
| 2020 | Age | Binary age | All | 0.0051 | -0.038–0.048 |
| | | | Females | -0.1136 | -0.176—0.051 |
| | | | Males | -0.4807 | -0.605—0.357 |
| | | Discrete age | All | 0.0032 | -0.031–0.037 |
| | | | Females | -0.0753 | -0.124—0.026 |
| | | | Males | -0.2837 | -0.364—0.203 |
| | | Continuous age | All | 0.0102 | -0.027–0.047 |
| | | | Females | 0.0657 | 0–0.132 |
| | | | Males | -0.2245 | -0.367—0.082 |
| | Residency status | Immigration status | All | 0.0631 | 0.019–0.107 |
| | | | Females | -0.0319 | -0.093–0.03 |
| | | | Males | 0.0025 | -0.157–0.162 |
| | | Settled status | All | -0.0184 | -0.061–0.025 |
| | | | Females | -0.1451 | -0.206—0.084 |
| | | | Males | -0.2971 | -0.435—0.159 |
| | Territory quality | Oak abundance | NA | 0.7269 | 0.705–0.749 |
| | | Average territory density | NA | 0.6088 | 0.586–0.632 |
| | | Edge distance index | NA | 0.7707 | 0.755–0.786 |
| | | Nest-box popularity index | NA | 0.1573 | 0.127–0.188 |
| | Reproductive | Clutch size | NA | -0.0643 | -0.099—0.03 |
| | | Number of chicks | NA | 0.0122 | -0.02–0.045 |
| | | Number of fledglings | NA | 0.1018 | 0.066–0.138 |
| | | Binary success | NA | 0.0570 | 0.018–0.096 |
| 2021 | Age | Binary age | All | -0.0688 | -0.102—0.036 |
| | | | Females | -0.0361 | -0.095–0.023 |
| | | | Males | 0.1431 | 0.056–0.23 |
| | | Discrete age | All | -0.0361 | -0.057—0.016 |
| | | | Females | 0.0380 | -0.004–0.08 |
| | | | Males | 0.1339 | 0.067–0.2 |
| | | Continuous age | All | -0.0315 | -0.07–0.007 |
| | | | Females | -0.0896 | -0.146—0.033 |
| | | | Males | 0.3546 | 0.151–0.558 |
| | Residency status | Immigration status | All | 0.0347 | 0.001–0.068 |
| | | | Females | 0.0551 | -0.006–0.116 |
| | | | Males | -0.1153 | -0.202—0.029 |
| | | Settled status | All | -0.0099 | -0.043–0.023 |
| | | | Females | 0.0707 | 0.011–0.13 |
| | | | Males | -0.0861 | -0.17—0.003 |
| | Territory quality | Oak abundance | NA | 0.7820 | 0.762–0.802 |
| | | Average territory density | NA | 0.6958 | 0.68–0.712 |
| | | Edge distance index | NA | 0.7398 | 0.726–0.754 |
| | | Nest-box popularity index | NA | 0.1864 | 0.152–0.220 |
| | Reproductive | Clutch size | NA | -0.0171 | -0.051–0.017 |
| | | Number of chicks | NA | 0.0712 | 0.036–0.107 |
| | | Number of fledglings | NA | 0.1712 | 0.137–0.205 |
| | | Binary success | NA | 0.2060 | 0.168–0.244 |
| 2022 | Age | Binary age | All | 0.0175 | -0.018–0.053 |
| | | | Females | 0.0849 | 0.024–0.146 |
| | | | Males | -0.1108 | -0.19—0.032 |
| | | Discrete age | All | 0.0319 | 0.008–0.056 |
| | | | Females | -0.0128 | -0.048–0.023 |
| | | | Males | -0.0476 | -0.092—0.003 |
| | | Continuous age | All | -0.0338 | -0.069–0.001 |

| Year | Focal trait type | Focal trait | Individuals | *r* | SE intervals |
|---|---|---|---|---|---|
| | | | Females | 0.1149 | 0.06–0.17 |
| | | | Males | -0.1203 | -0.19—0.05 |
| | Residency status | Immigration status | All | -0.0328 | -0.068–0.002 |
| | | | Females | -0.0316 | -0.09–0.026 |
| | | | Males | -0.0993 | -0.178—0.02 |
| | | Settled status | All | 0.1053 | 0.068–0.142 |
| | | | Females | 0.0185 | -0.045–0.082 |
| | | | Males | 0.2716 | 0.193–0.35 |
| | Territory quality | Oak abundance | NA | 0.7450 | 0.724–0.766 |
| | | Average territory density | NA | 0.5982 | 0.565–0.631 |
| | | Edge distance index | NA | 0.7330 | 0.718–0.748 |
| | | Nest-box popularity index | NA | 0.2888 | 0.258–0.319 |
| | Reproductive | Clutch size | NA | 0.1064 | 0.065–0.148 |
| | | Number of chicks | NA | 0.0889 | 0.05–0.128 |
| | | Number of fledglings | NA | 0.1425 | 0.103–0.181 |
| | | Binary success | NA | 0.1152 | 0.072–0.159 |

Table S5 – Temporal repeatability of age composition, residency status composition, average territory quality and average reproductive output of neighbourhoods at spatial scales of 25, 50, 100, 150 and 200ha. Repeatability was derived from the intra-class correlation coefficient (ICC) from a linear mixed-effects model, where the ratio index of within versus outside the defined spatial scale of the average measure was the response variable, and year and spatial scale area categorical grouping variables.

| | **Repeatability at spatial scales:** | | | | |
|---|---|---|---|---|---|
| **Trait** | **25ha** | **50ha** | **100ha** | **150ha** | **200ha** |
| Proportion adults | 0.036 | 0.050 | 0.057 | 0.043 | 0.031 |
| Proportion local recruits | 0.076 | 0.102 | 0.135 | 0.157 | 0.173 |
| Proportion settled individuals | 0.024 | 0.035 | 0.048 | 0.057 | 0.062 |
| Mean oak abundance | 0.971 | 0.973 | 0.969 | 0.962 | 0.955 |
| Mean territory density | 0.968 | 0.974 | 0.976 | 0.975 | 0.975 |
| Mean EDI | 0.965 | 0.965 | 0.967 | 0.970 | 0.973 |
| Mean box popularity index | 0.916 | 0.939 | 0.959 | 0.967 | 0.971 |
| Mean clutch size | 0.234 | 0.294 | 0.364 | 0.412 | 0.447 |
| Mean chick number | 0.090 | 0.118 | 0.165 | 0.194 | 0.219 |
| Mean fledgling number | 0.182 | 0.240 | 0.314 | 0.361 | 0.398 |
| Proportion binary successes | 0.084 | 0.111 | 0.150 | 0.176 | 0.202 |

———. 1977. Song and territory in the great tit Parus major. in B. Stonehouse and C. M. Perrins, eds. Evolutionary Ecology. MacMillan, London.

———. 1982. Territorial Defence in the Great Tit (Parus major): Do Residents Always Win? Behavioral Ecology and Sociobiology 11:185–194.

Paton, P. W. C. 1994. Effect of edge on avian nest success: how strong is the evidence? Conservation Biology 8:17–26.

Sandell, M., and H. G. Smith. 1991. Dominance, prior occupancy, and winter residency in the great tit (Parus major). Behavioral Ecology and Sociobiology 29:147–152.

Slagsvold, T., and K. L. Wiebe. 2018. Immigrants and locally recruited birds differ in prey delivered to their offspring in blue tits and great tits. Animal Behaviour 139:127–135.

Verhulst, S., C. M. Perrins, and R. Riddington. 1997. Natal dispersal of Great Tits in a patchy environment. Ecology 78:864–872.

Wilkin, T. A., D. Garant, A. G. Gosler, and B. C. Sheldon. 2006. Density effects on life-history traits in a wild population of the great tit Parus major: Analyses of long-term data with GIS techniques. Journal of Animal Ecology 75:604–615.

———. 2007a. Edge effects in the great tit: Analyses of long-term data with GIS techniques. Conservation Biology 21:1207–1217.

Wilkin, T. A., C. M. Perrins, and B. C. Sheldon. 2007b. The use of GIS in estimating spatial variation in habitat quality: A case study of lay-date in the Great Tit Parus major. Ibis 149:110–118.
